# Systematic Review of Over A Century of Global Bioscience Research

**DOI:** 10.1101/2025.10.14.682434

**Authors:** Okechukwu Kalu Iroha, Dauda Wadzani Palnam, Peter Abraham, Israel Ogwuche Ogra, Ndukwe K. Johnson, Elkanah Glen, Dasoem Naanswan Joseph, Seun Cecilia Joshua, Grace Peter Wabba, Morumda Daji, Dogara Elisha Tumba, Mercy Nathaniel, Emohchonne Utos Jonathan, Samson Usman, Mela Ilu Luka, Vaibhav B. Sabale, Emmanuel Oluwadare Balogun, Umezuruike Linus Opara

**Author notes:** Corresponding author: **Dauda Wadzani Palnam,**.

## Abstract

Bioscience encompasses studies on living organisms, their components, and their interactions, with the main aim of translating research into useful applications for medical, clinical, industrial, and environmental uses. The broad field of bioscience has witnessed tremendous growth in research output. This study was designed to evaluate the current trends in bioscience research using bibliometric approaches and develop future policy pointers following the innovative systems framework. Research articles focusing on bioscience published between 1883 and 2024 were sourced from the Scopus database. From the retrieved dataset, relevant articles were systematically screened and included for the analysis. Bibliometrix package, VOSviewer, and Microsoft Excel were used to analyze and visualize the parameters. 8,678 documents involving 30,380 authors from diverse institutions across the globe were published in 3,549 sources, with an annual rise of 4.41%. Exponential growth in bioscience research output was observed after 2007 and remained steady till 2024. The United States and China were leading nations for bioscience research. The most relevant affiliation was the University of California, United States, while the most relevant source was the *Brazilian Journal of Medical and Biological Research*, in terms of the number of publications. Among the authors, Wang Y had the highest number of documents, while the most impactful author was Zhang J. The most frequent keywords in bioscience research included *biological research*, *genetics*, *procedures*, *metabolism*, *biology*, *DNA*, *gene expression*, *proteins*, *genomics*, and *fluorescence*. Furthermore, themes such as *bioinformatics*, *microRNA*, *synthetic biology, CRISPR/Cas9*, *deep learning*, *multi-omics*, and *big data* represented the recent research interests. The bioscience research landscape is defined by the dominance of high-income countries and a shift toward omics-driven and data-intensive approaches. Yet global participation remains uneven, with limited representation from low- and middle-income regions. This study highlights the need for inclusive, globally coordinated strategies that foster equitable access, capacity building, and cross-regional collaboration.

## INTRODUCTION

The international science system is made up of instantly shared knowledge and expertise in the universal language of scientific study and mutually beneficial interactions among scholars in multidisciplinary relationships, all of which are facilitated by fundamental factors such as funds, facilities, tools, regulations, and education and training (Marginson, 2022). As an important aspect of science, bioscience developments are crucial to illness prevention, environmental protection, and economic prosperity (Hamburg et al., 2022). Understanding the biological underpinnings of health through the fundamental fields of physiology, biochemistry, microbiology and pathology, pharmacology, immunology, neuroscience, and genetics is made easier with the help of bioscience expertise (McColl et al., 2012). Accordingly, Flier (2017) opined that clinical investigation and translational research that connects basic sciences to medical use, in a range of disciplines, are all included in the wide category of bioscience research. Thus, the broad bioscience term encompasses studies on living organisms, their components, and their interactions with the environment. The array of bioscience publications has grown significantly since about 20 years, with many of them occupying specialized niches that represent emerging fields and technology (Kilkenny et al., 2012).

The field of bioscience has grown tremendously. The ability of students and researchers to perform biological and clinical tests, observe treatment effects, and report findings effectively depends on their understanding of bioscience (Berre et al., 2024). Bioscience is taught using a variety of methods, and allied fields are increasingly using digital tools (Manchester and Roberts, 2024). New opportunities for enhancing learning are presented by the emergence of virtual environments and modeling, as well as the usage of educational platforms on the web (Kaltsidis et al., 2021; Higgins et al., 2024).

The innovation systems framework can be used to understand the growth of the bioscience research and education landscape. In general, the theory of innovation systems seeks to comprehend how a collection of organizations, structures, systems, and people can collaborate to stimulate growth in a specific national, regional, or specialized location, or related to the advancement of a modern technology (Lundvall, 2007; Touzard et al., 2015). Innovation systems influence bioscience research and education by promoting technical improvements, multidisciplinary cooperation, and knowledge diffusion. Research institutions are examples of innovation ecosystems providing financing and support for major bioscience advancements such as omics and individualized healthcare (Bradke et al., 2023). Also, creative learning incorporates artificial intelligence, digital tools, and student-oriented educational efforts to enhance biology education by rendering it more experiential (Bhardwaj et al., 2022; Kim et al., 2023). In addition, multi-sector alliances boost bioscience research by bringing together universities, industry, and governmental organizations to speed breakthroughs and commercialization (Altshul et al., 2019). Thus, innovation systems, which include institutions, regulations, collaborations, and economic factors, have an impact on bioscience research and education by influencing how knowledge is created, implemented, and appraised.

Literature reviews are essential in academic research to determine the current state of a topic (Linnenluecke et al., 2020). In order to evaluate the expansion and impact of a subject area, several disciplines have begun to employ better, more organized methods that emphasize quantitative approaches (Keathley-Herring et al., 2016). Bibliometric evaluation is a popular and accurate method for searching through and assessing vast amounts of scientific knowledge (Donthu et al., 2021). Bibliometric techniques shed light on fresh directions in a field while assisting in comprehending the finer points of a certain topic’s development (Linnenluecke et al., 2020). This emphasizes why it was chosen for this inquiry.

Despite the innovations in bioscience research and education, there are not many bibliometric reviews of the field. Abdullah (2022) analyzed the trends in biology education, while Barbosa and Galembeck (2022) evaluated the research landscape of biochemistry education. Both present a narrow focus on the wide field of bioscience, concentrating on biology and biochemistry, respectively. Jang and Kim (2014) analyzed the research output of science, technology, and bioscience, albeit with a focus only on the Asian continent. Thus, there is a lack of bibliometric reviews that particularly evaluate the broad scope of bioscience research and education. These reviews are useful for determining past research patterns and new subjects, evaluating research partnerships and output, measuring the impact of research, filling in knowledge gaps, and determining the direction of future studies. Therefore, the purpose of this study is to use bibliometric techniques to examine the research output on bioscience research and education. This review covers 141 years (1883-2024), delivering a long-term perspective hardly found in bioscience bibliometric analyses. It captures the transition from early scientific methods to modern research trends. Also, it expands the conventional publication evaluation by merging bibliometrics with innovation systems theory. This places scientific research within the contexts of technological growth, institutional transformation, and regional innovation plans, connecting insights based on data to policy significance, as conceptualized in the schematic diagram in Figure 1.

**Figure 1:**
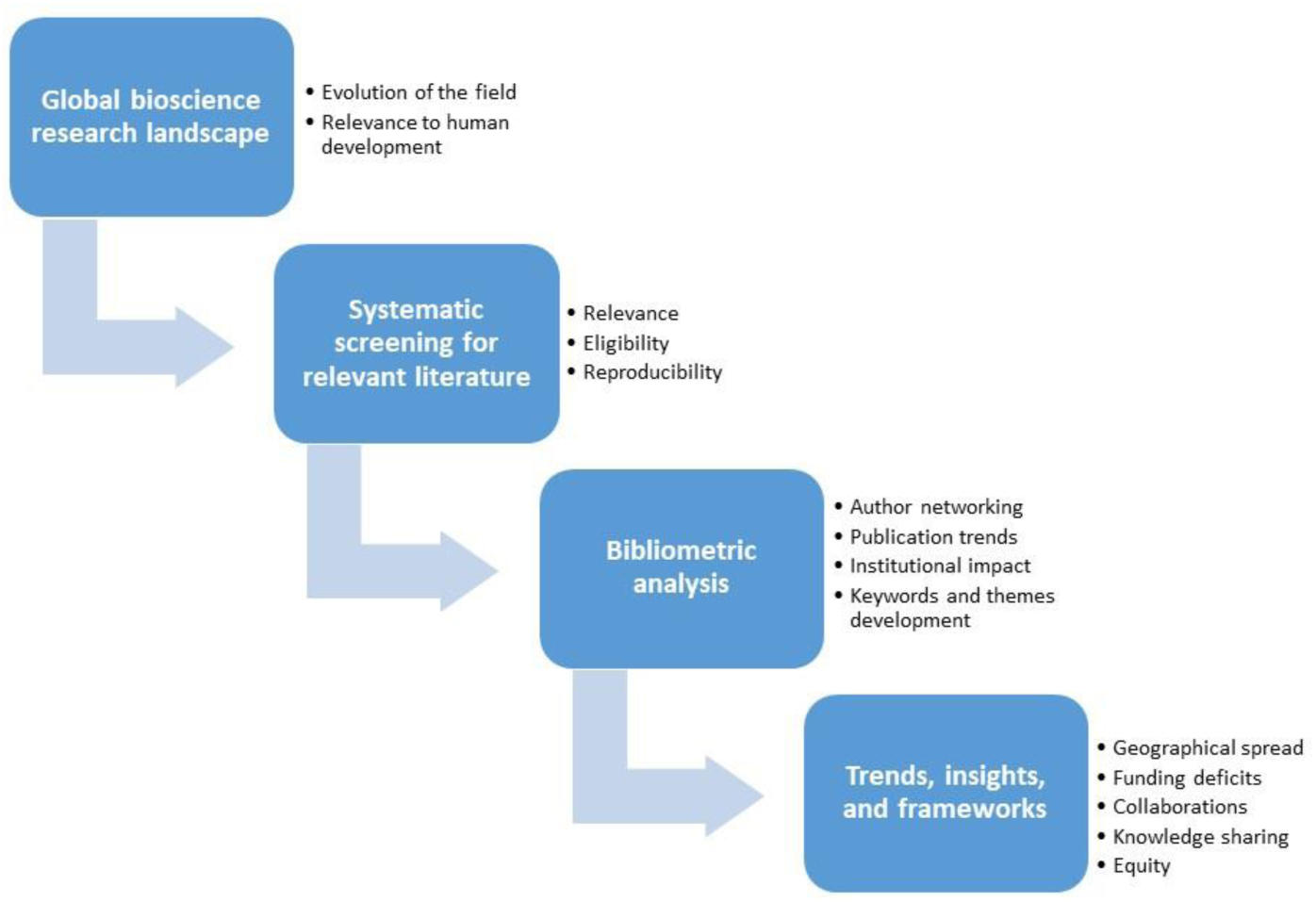
Schematic diagram conceptualizing the research

To achieve this aim, the review used the following research objectives: identify the general trends in bioscience research and education, evaluate country performance and global collaborations, evaluate the contributions of affiliations, sources, and authors, analyze the evolution of keywords and author-suggested keywords, and identify the most cited documents.

The arrangement of this investigation is presented thus. In section 2, the data collection and analysis approaches are provided. In section 3, the findings generated from the review are presented, which involve the publication statistics, highly-cited articles, authors, institutions, and sources/journals, geographical evaluation of documents, and keywords analysis. Section 4 offers fundamental discourse and policy proposals on bioscience research, highlighting the study’s boundaries. Section 5 concludes the review by presenting a recap of major findings.

## METHODOLOGY

### Data collection

The data sets used for this analysis were sourced and retrieved from the Scopus database on 25/8/2024, at 7:45 pm. This database is widely recognized for its coverage and reliable content, housing numerous publications from reputable publishers. The period was set at 1883 to 2024 to cover the onset of research output on the subject.

#### Search strategy

The search was performed on the titles, abstract and keyword fields using the search string (("bioscience research") OR ("biological research") OR ("biological science research") OR ("natural science research") OR ("life sciences education”) OR ("life sciences research”) OR ("bioscience education") OR ("bioscience analysis") OR ("bioscience exploration") OR ("bioscience investigation") OR ("bioscience education") OR ("biological science education") OR ("bioscience learning") OR ("undergraduate bioscience") OR ("higher education bioscience") OR ("bioscience academics") OR ("bioscience training") OR ("bioscience teaching") OR ("bioscience curriculum") OR ("bioscience pedagogy") OR ("bioscience classroom")).

#### Screening process

A total of 12,803 results were retrieved. However, 8,773 documents were remaining after language (English) and document type filtering (Research and Review articles) were performed (4,030 documents were removed at this stage). The search results were exported as a CSV file for screening. The retrieved data were manually screened to systematically exclude publications that are not in line with the subject area. The PRISMA (Preferred Reporting Items for Systematic Reviews and Meta-Analyses) guidelines were used for title and abstract screening, and full text review, where necessary, following the exclusion criteria (Figure 2). The inclusion criteria were that the accepted articles must be research or review papers, published in English, and at the final stages of publication. Documents such as textbooks, handbooks, commentaries, and articles not published in English, or articles retrieved in the initial publication stages, were excluded. All co-authors agreed to follow a common protocol (PRISMA) and apply consistent screening criteria. Disagreements were handled through discussion or consensus gatherings. During the manual screening of titles and abstracts, 95 documents were removed as they were not in line with the topic. A total of 8,678 documents were selected and included as an exact representation of relevant documents covering research on the search strings. This set of data was used for further analysis.

**Figure 2:**
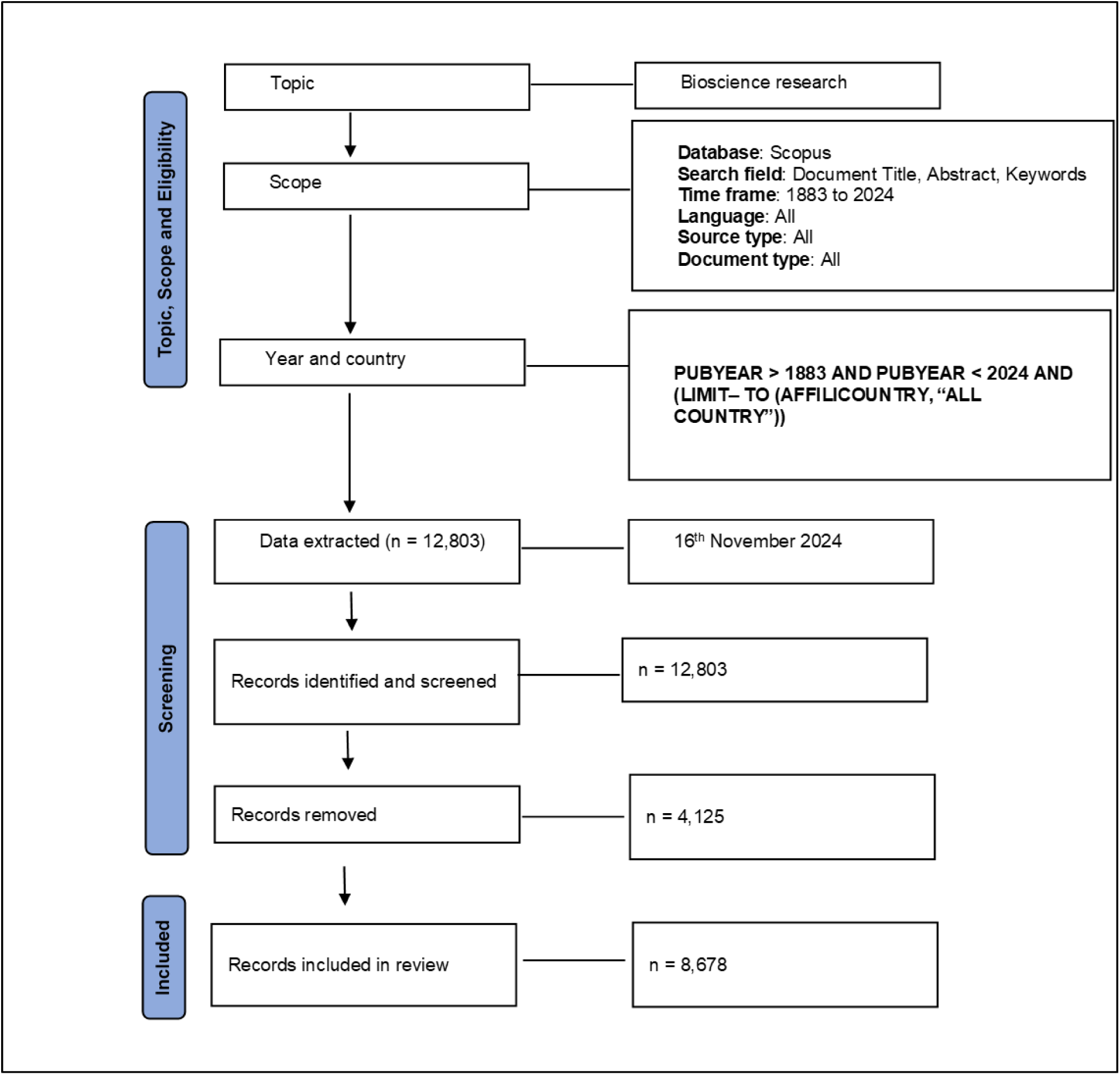
Flowchart for the retrieval, screening and inclusion of analyzed studies

#### Data analysis

The raw search results from the Scopus database were exported as a CSV file for bibliometric analysis. The Bibliometrix software (Aria and Cuccurullo, 2017) and VOSviewer software (Van Eck and Waltman, 2010) were used for the bibliometric analyses. The Bibliometrix software enabled the analysis of global trends in productivity, impact, and conceptual structure. Network maps for keyword co-occurrence and country co-authorship were generated with the VOSviewer software. Microsoft Excel was used for visualizing the data analysis with clustered bar charts. For the synthesis of insights, both bibliometric and systematic review approaches were integrated. This involved the analysis of the most cited documents to generate key themes and concepts.

## RESULTS

### Analysis of general trends in bioscience research and education

The details of the retrieved dataset are presented in Figure 3. The search for scientific information on bioscience in the Scopus database revealed that 8,678 documents were published in 3,549 sources between 1883 and 2024. These publications were contributed by 30,380 authors affiliated with diverse institutions across the globe. With an annual rise (compound annual growth rate) of 4.41%, there is a considerable increase in the number of publications on bioscience.

**Figure 3:**
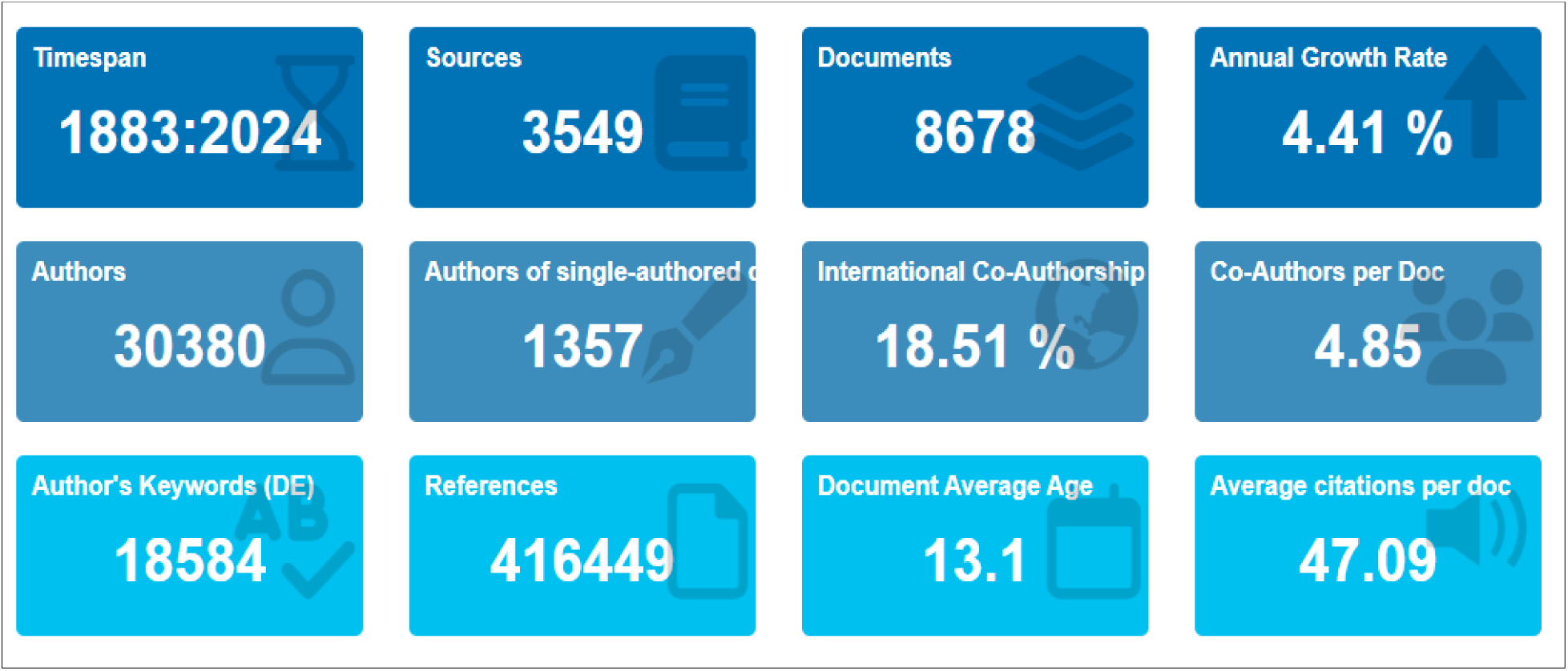
Summary of information on retrieved data for bioscience research and education

As shown in the chart in Figure 4, the publication production over time shows a period of inactivity between 1883 and 1950. However, from 1955, a lag phase of growth in bioscience research activity started and extended to 1990. There was an increased research output on bioscience in the 1990s, representing an initial steady growth until 2007. Exponential growth in bioscience research output was observed after 2007 and remained steady till 2024. The highest average total citations of bioscience research and education articles per year were observed in 2002 and 2014, as shown in Figure 5. Figure 6 depicts the bioscience research document distribution according to type, showing that of the 8,678 documents, research articles were the most abundant documents, totaling 6,795, while reviews were 1,883.

**Figure 4:**
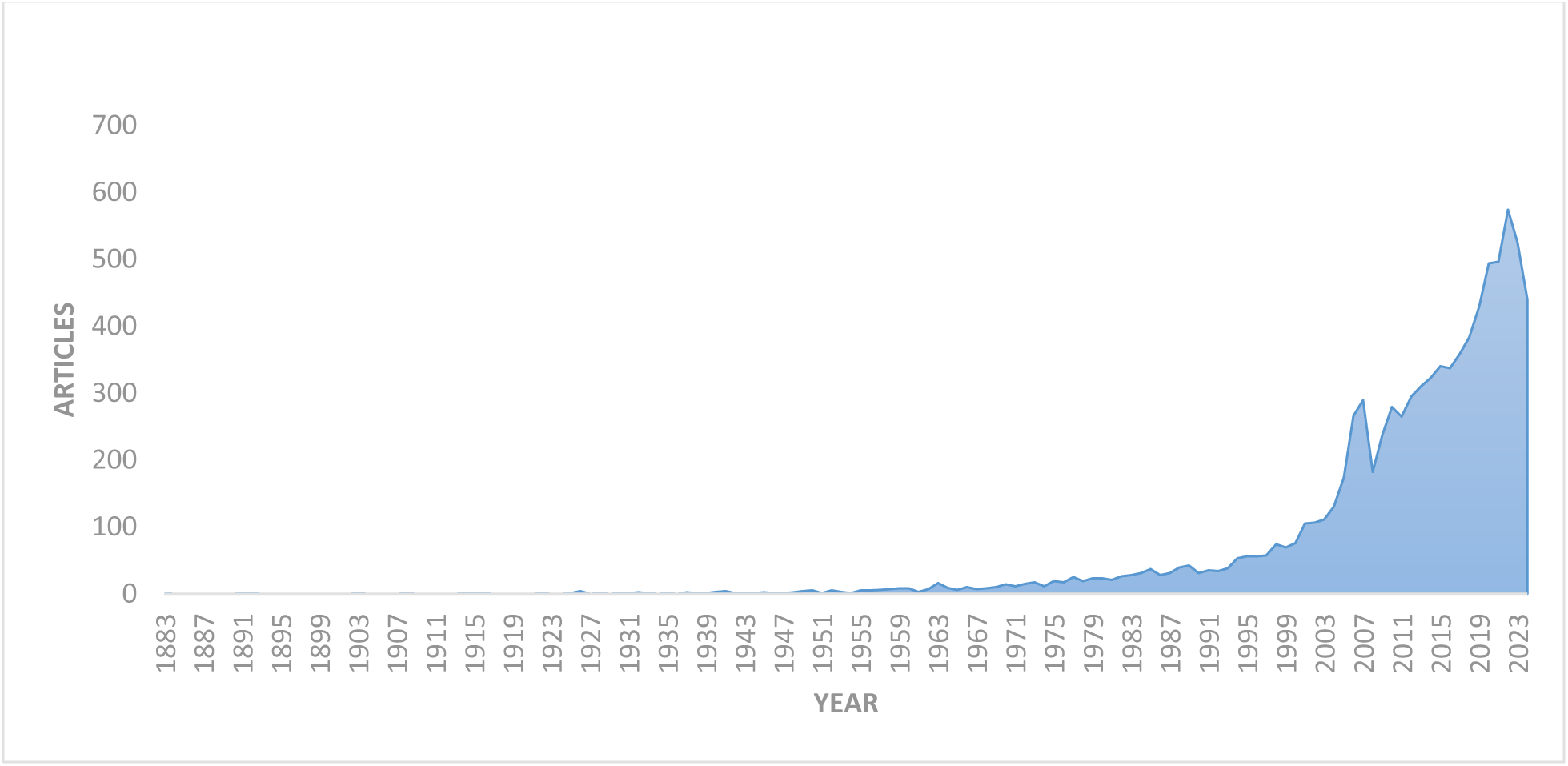
Annual publications production on bioscience research

**Figure 5:**
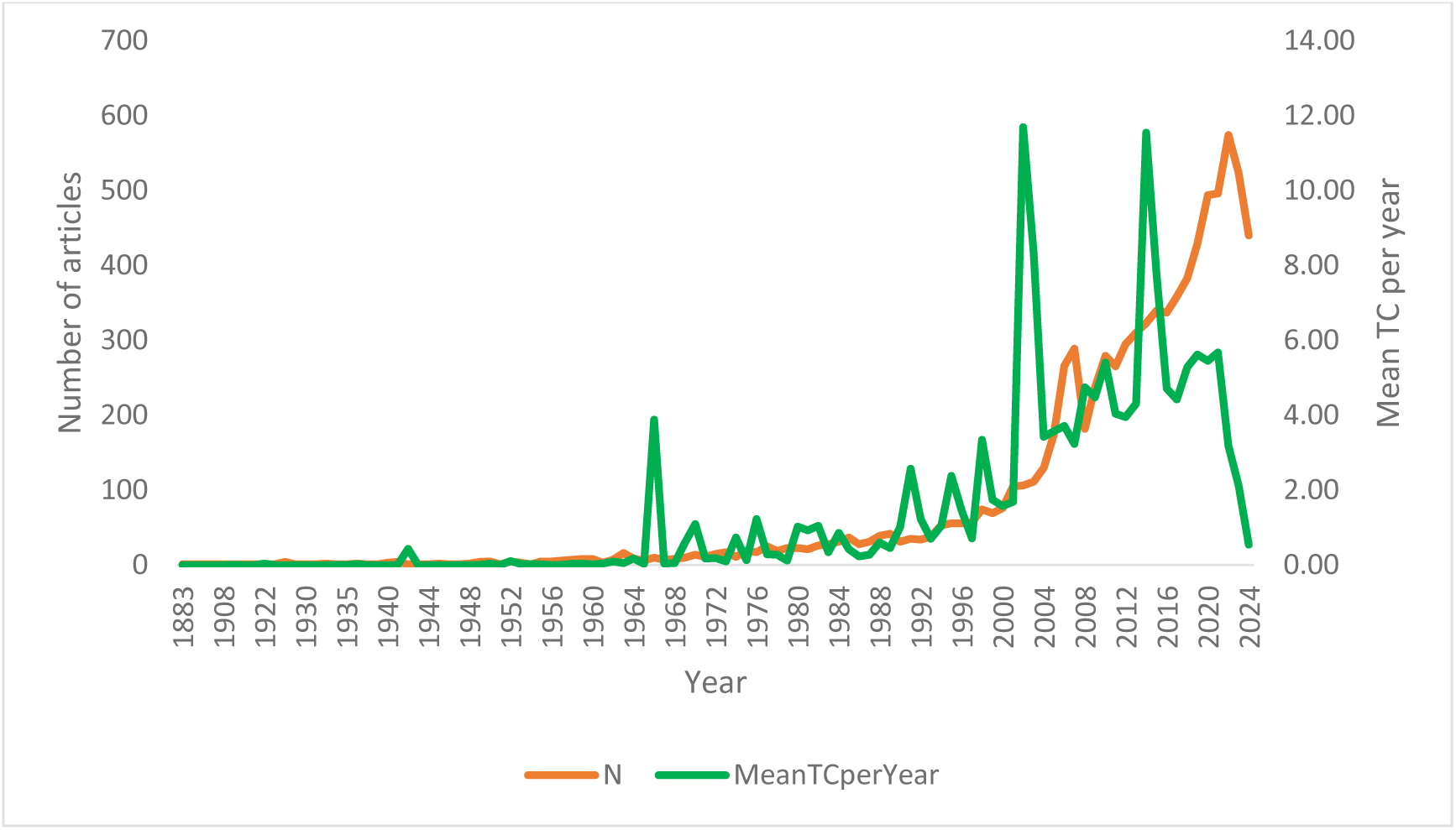
Comparison of yearly scientific production and citation

**Figure 6:**
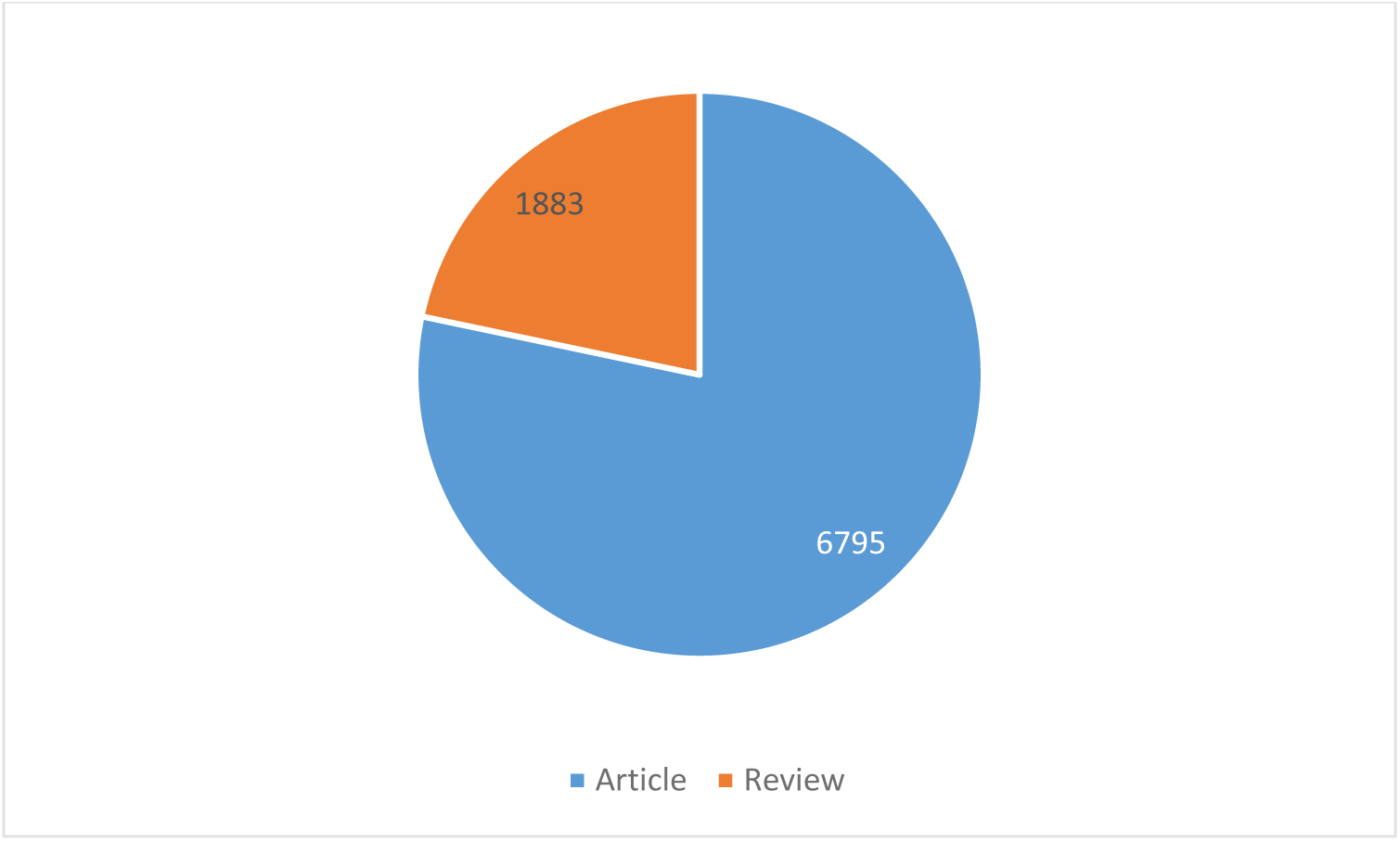
Document distribution by type

#### Analysis of country performance and collaborations

The map in Figure 7A highlights the worldwide distribution of publications on bioscience. The deep blue reflects countries and regions where there are a high number of publications. From the map, the United States and China are the countries with the highest research output on bioscience. The distribution of documents across countries, as shown in Figure 7B, further corroborates that the United States and China are leading nations in bioscience research. They are followed by other developed nations such as the United Kingdom, Japan, Germany, and Brazil.

**Figure 7A:**
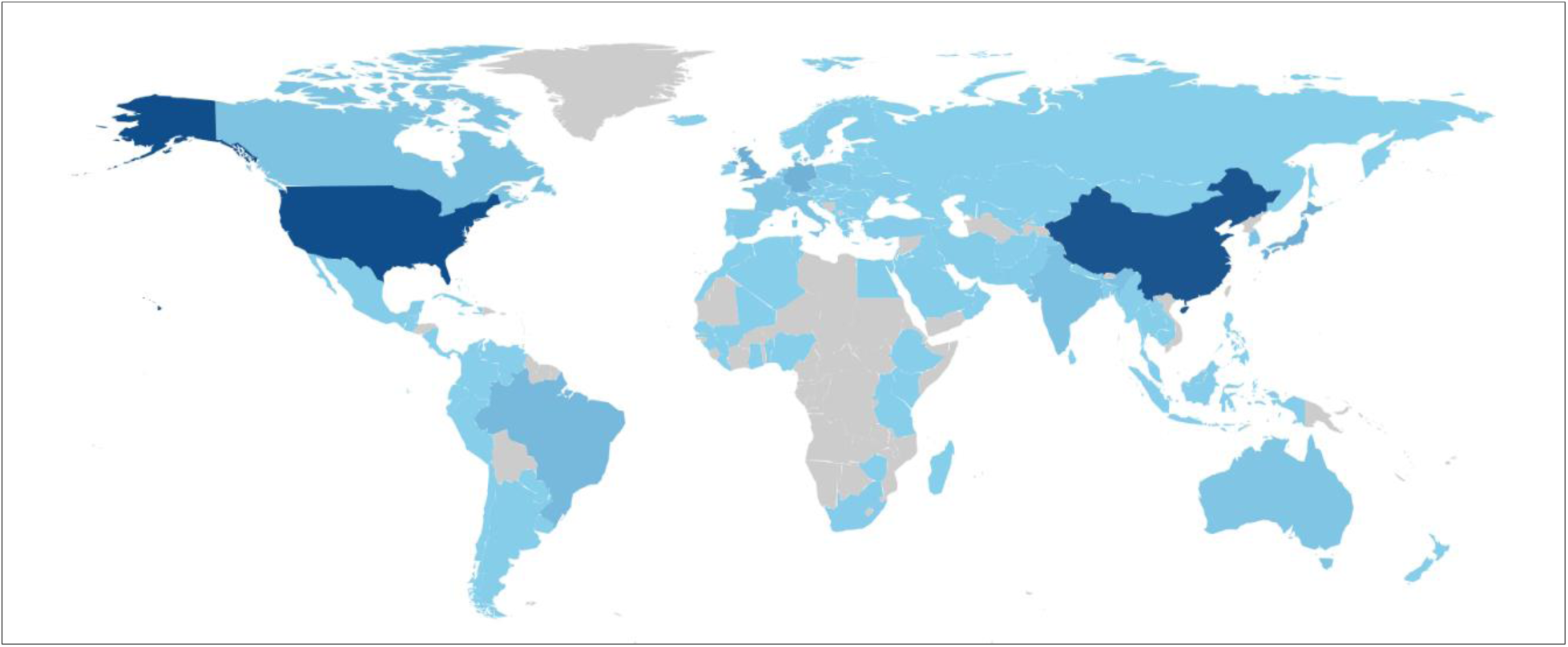
Global distribution of publications on bioscience research

**Figure 7B:**
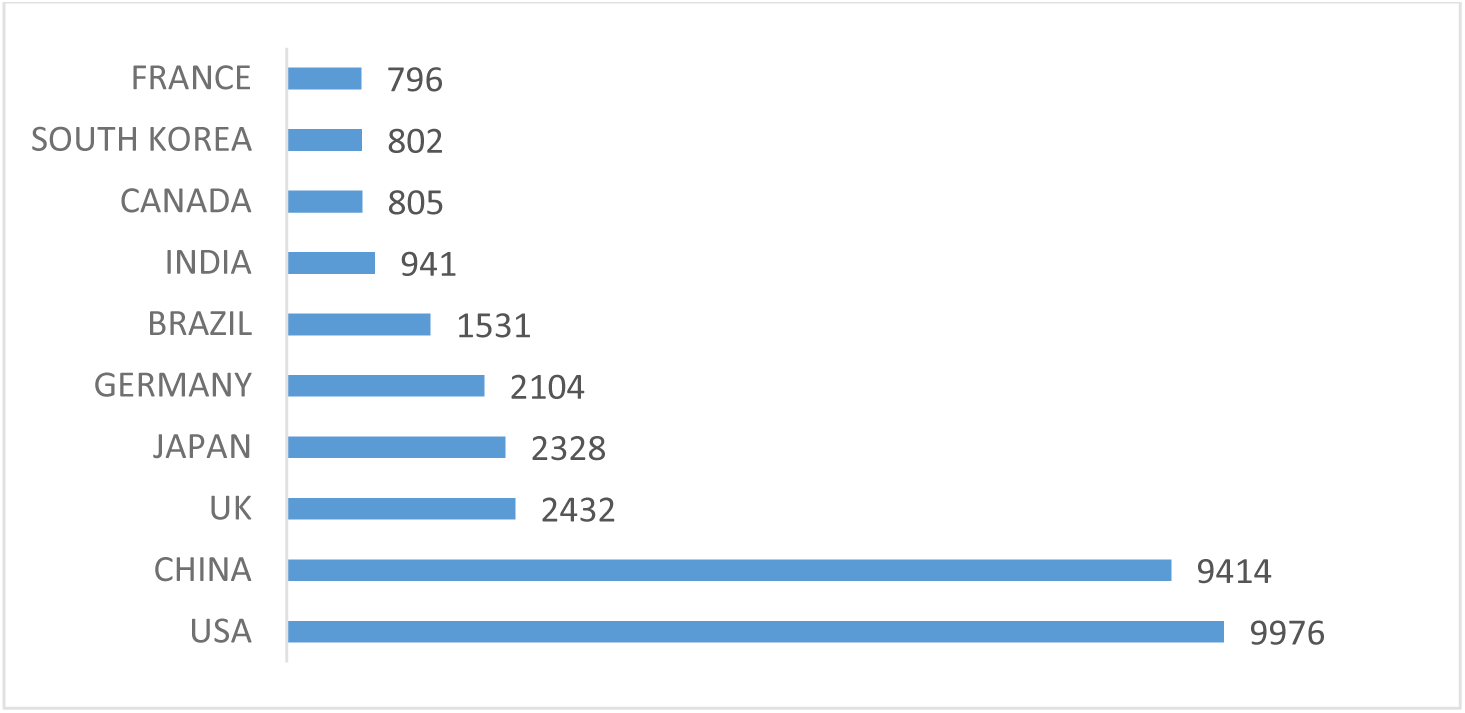
Distribution of documents/articles across top 10 countries

The analysis of country co-authorship using VOSviewer revealed that out of 242 countries, 74 met the threshold of at least 5 documents per country, further suggesting a global advancement of the biosciences. The network map in Figure 8 depicts the co-authorship collaborations among countries. Each node in the map represents a country, while the weight of each node denotes the number of documents for that country. Each line in the map indicates a connection between countries. Eight (8) clusters represented with colours were identified. From the network map, the United States, China, Germany, Japan, India, and Brazil had a higher number of documents. Also, in Table 1, research hotspots such as the United States, the United Kingdom, and Germany showed higher linkages in relation to other countries.

**Figure 8:**
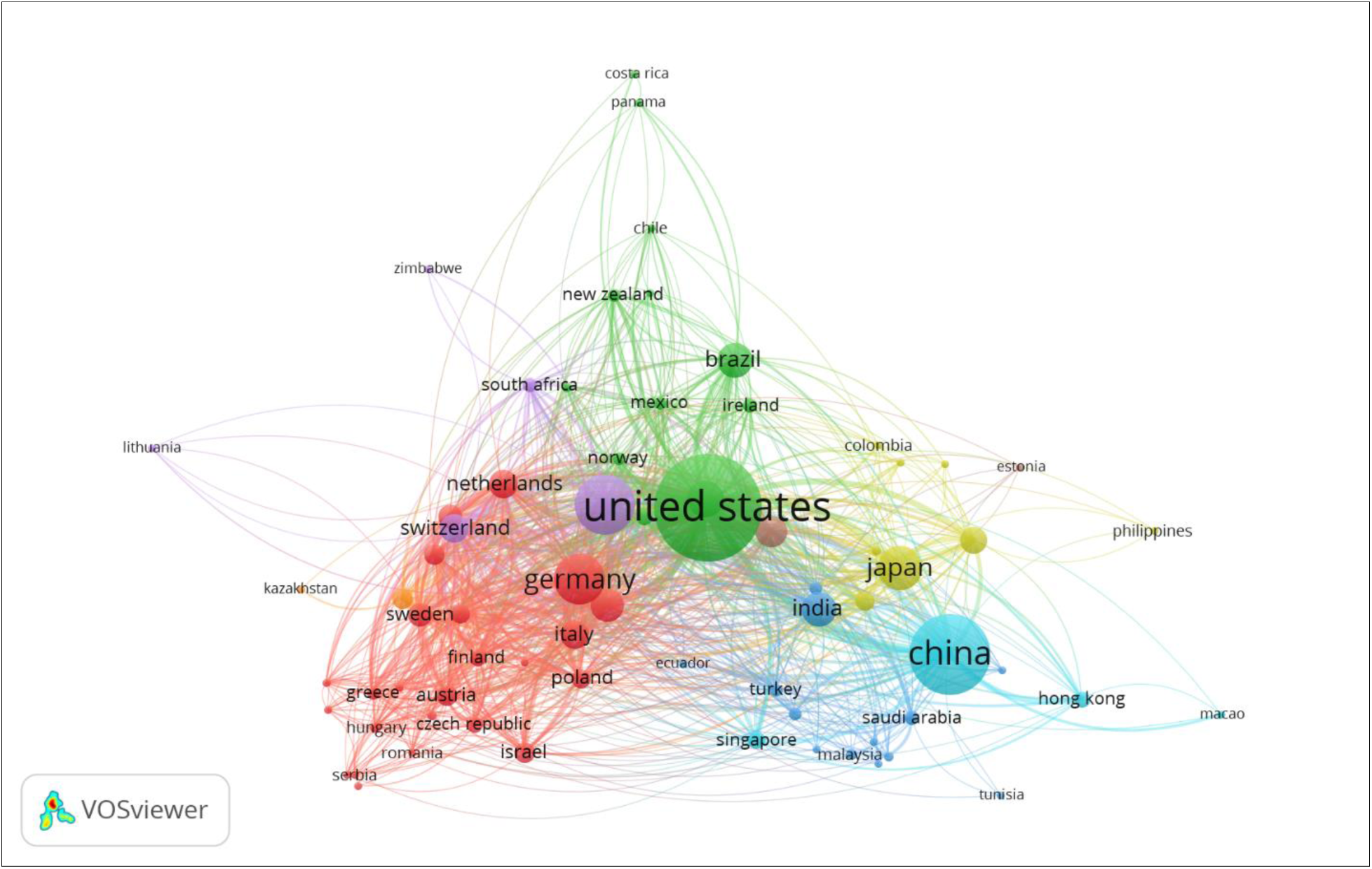
Country co-authorship network based on number of documents

**Table 1:**
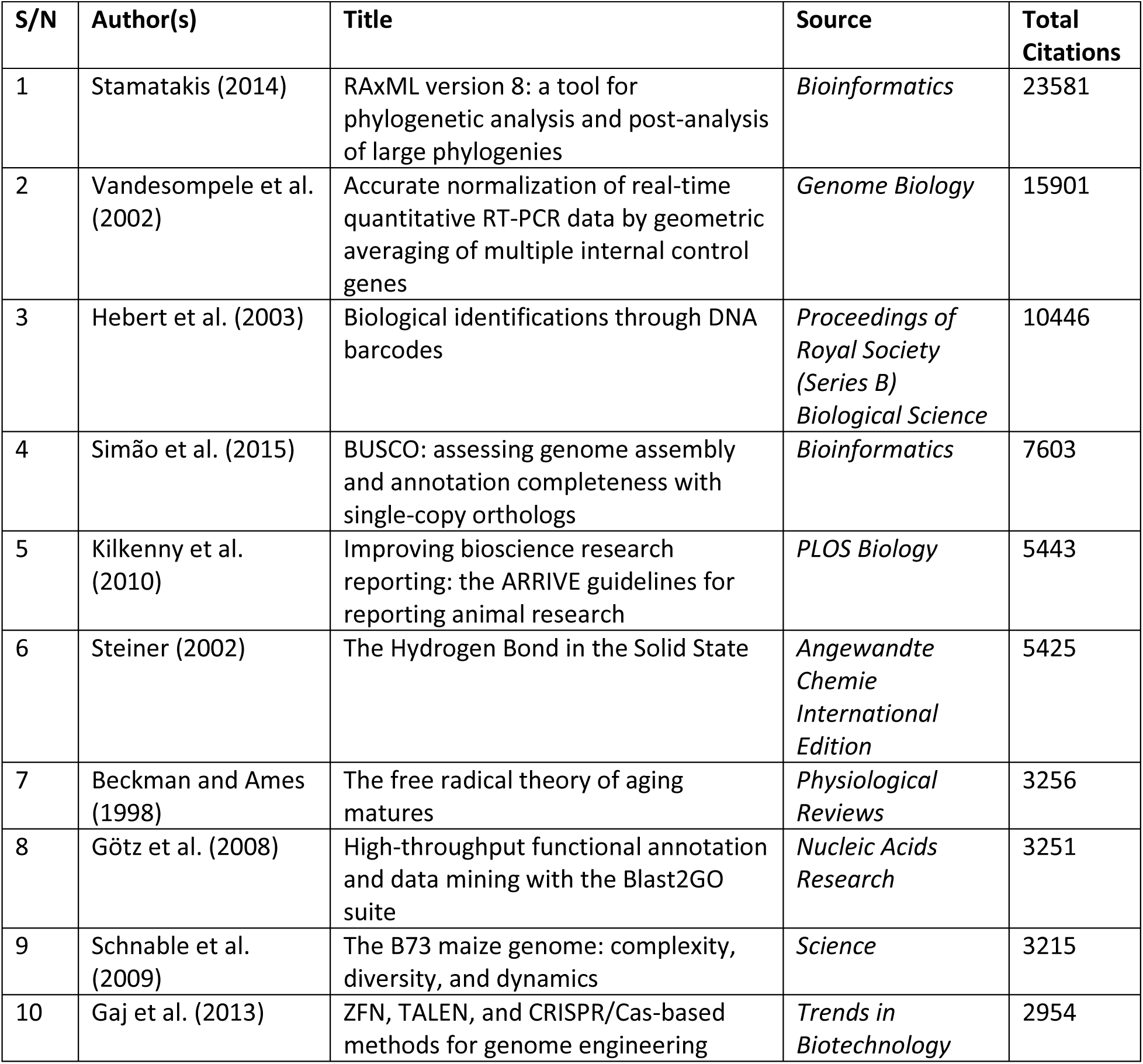
Details of the top 10 most cited documents

Figure 9 reflects the global distribution of single and multiple country collaborations for publications in bioscience research and education. A surprising observation from the chart is that all the top 20 countries had more single-country publications than multiple-country publications.

**Figure 9:**
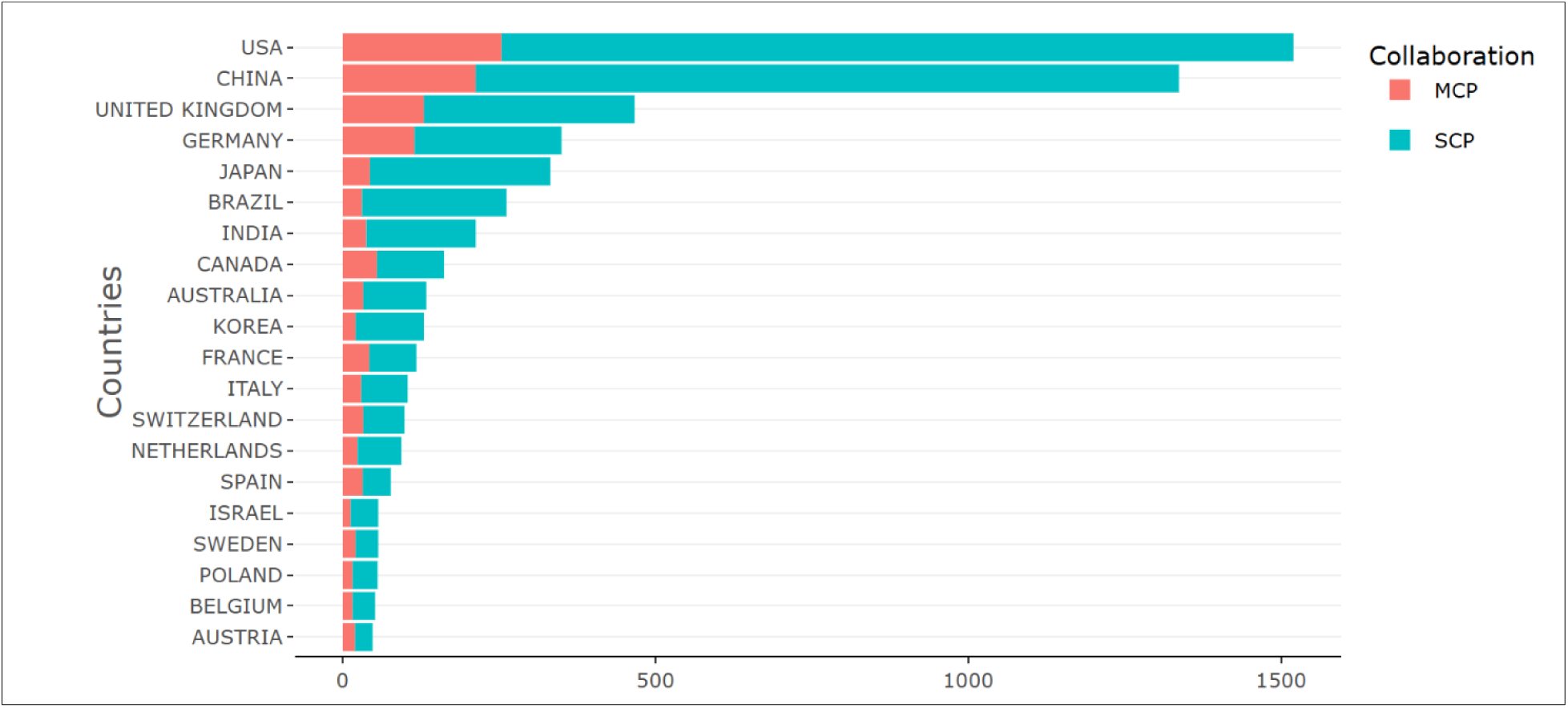
Global distribution of single and multiple country collaborations

As illustrated in Figure 10, the most cited country on bioscience research and education was the United States, with a total of 96,817 citations. Germany, China, and the United Kingdom follow with 50,202, 32,630, and 29,470 citations, respectively. Also, among the highly cited countries are Belgium, Canada, and Switzerland.

**Figure 10:**
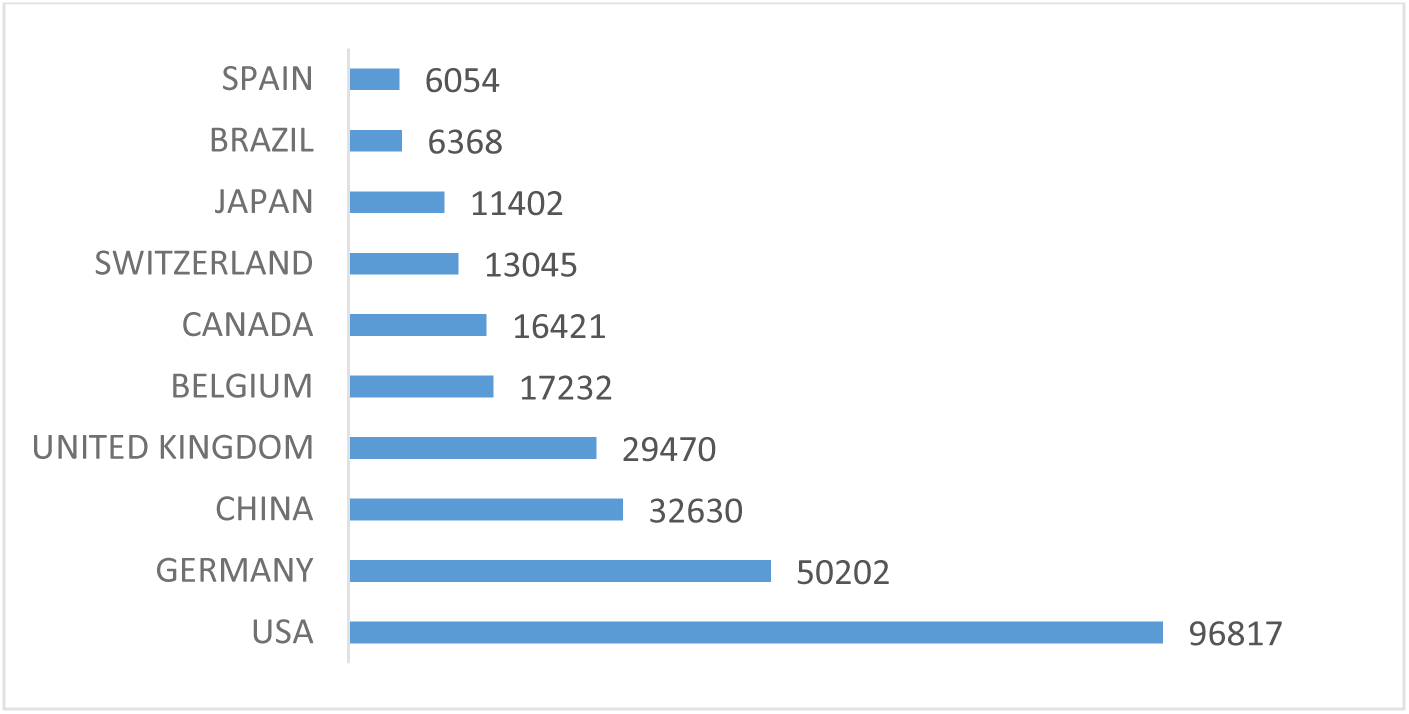
Top 10 most cited countries

#### Analysis of affiliations, sources, and authors’ contributions

From the analysis on the retrieved data, the institution with the most publications on the subject matter was the University of California, United States, with a total of 398 documents, as highlighted in Figure 11. This is followed by Universidade de São Paulo (349 documents) and University of Zurich (207). Others include Universidade Federal de São Paulo (175), Zhejiang University (172), and Tsinghua University (153 documents).

**Figure 11:**
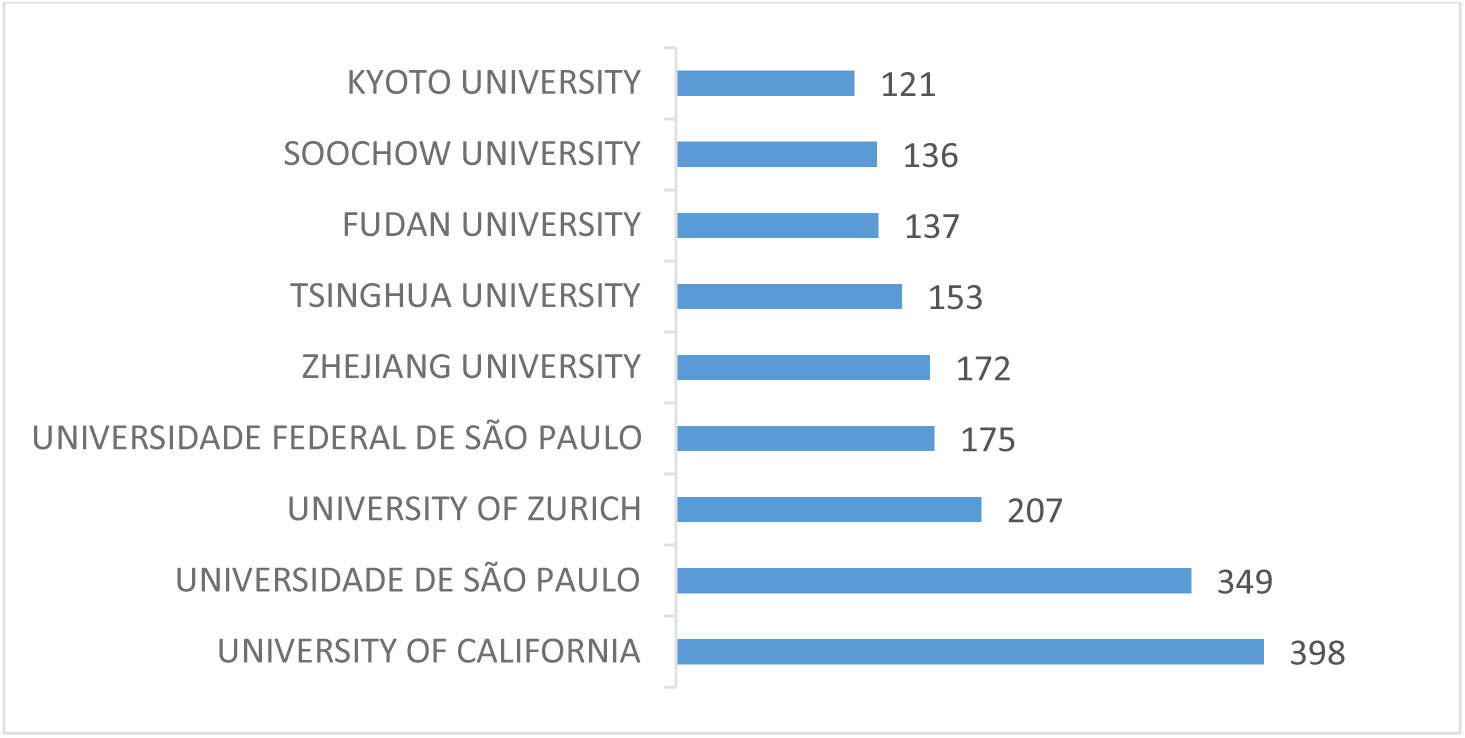
Institutions with the most publications on bioscience research

Figure 12 shows the top 10 most relevant sources out of the 3,549 sources on bioscience research, ranked according to the number of documents. The most relevant source in which bioscience-related research was published, with a total of 238 documents (25.2%), was the *Brazilian Journal of Medical and Biological Research*, published by Associação Brasileira de Divulgação Científica (Brazilian Association of Scientific Dissemination). Other relevant sources included *PLOS ONE* (123 documents, 13.04%), *Analytical Chemistry* (81 documents, 8.6%), *Nucleic Acids Research* (80 documents, 8.5%), *BMC Bioinformatics* (76 documents, 8.1%), and *Scientific Reports* (76 documents, 8.1%).

**Figure 12:**
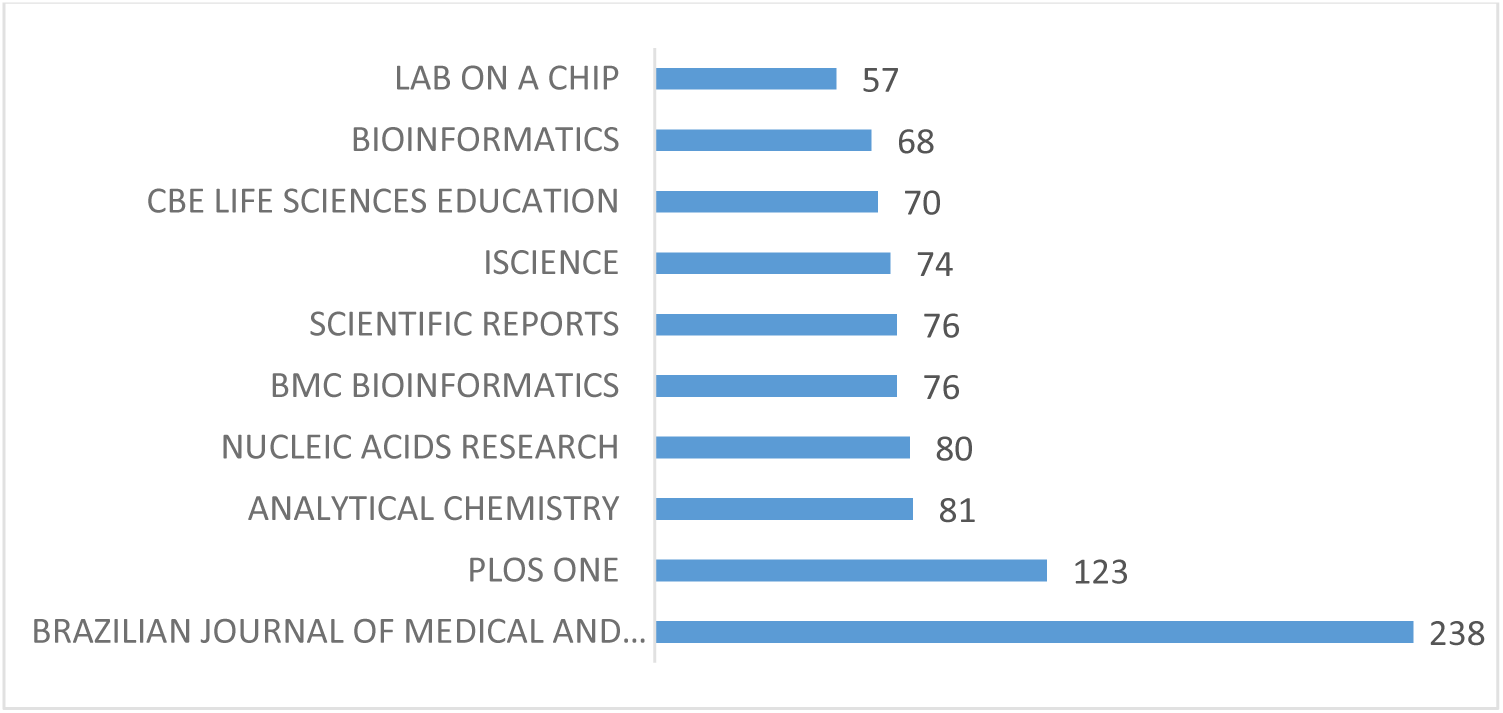
Top 10 relevant sources of research on bioscience

The network map of the sources as analyzed with VOSviewer showed that out of the 3,549 sources, 320 met the threshold of at least 5 documents. Twenty-two (22) clusters with different counts of items (sources) were also identified. The overlay visualization map is shown in Figure 13. Each node represented a source, while the size of the node depicted the number of documents. *Nature* and *Science* were the dominant sources before 2005. Relevant sources dominating between 2010 and 2017 included *Journal of Microbiology*, *BMC Bioinformatics*, *Bioinformatics*, *CBE Life Sciences*, *Lab on a Chip*, *Analytical Chemistry*, and *Analytical and Bioanalytical Chemistry*. However, Sc*ientific Reports*, *ACS Synthetic Biology*, *Frontiers in Bioengineering and Biotechnology*, *Genes*, *iScience*, *International Journal of Molecular Sciences*, *Frontiers in Genetics*, and *Heliyon* showed dominance from 2018 to 2022.

**Figure 13:**
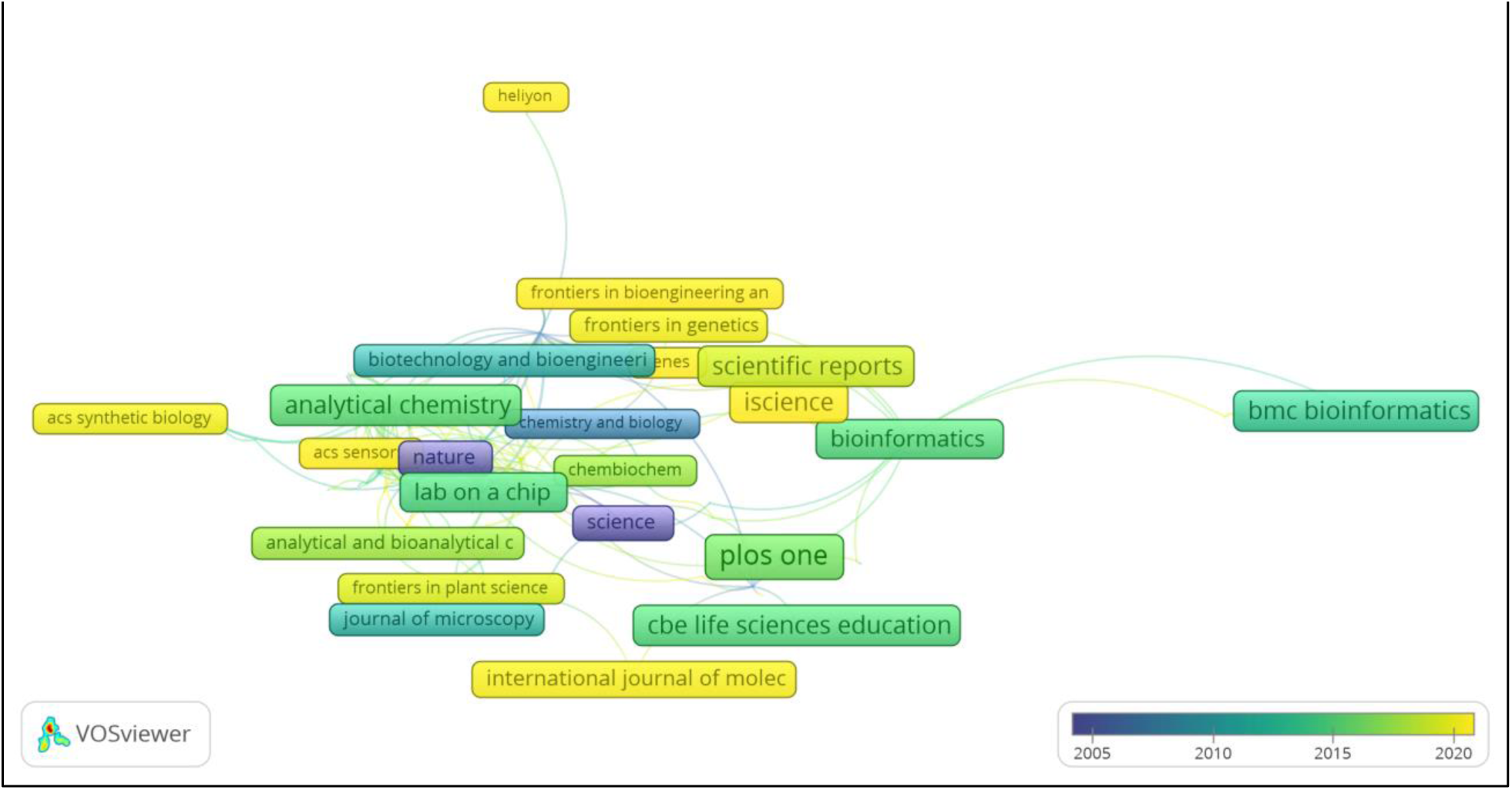
Sources/ (journals) network map

Within the reviewed period, 30,380 authors contributed to publications on bioscience research and education, with 1,357 (representing 4.47%) of them producing single-authored documents. The topmost producing authors according to the number of documents are presented in Figure 14. Wang Y had the highest documents with 128 publications. Zhang Y and Liu Y follow with 119 and 105 documents, respectively. The other relevant authors and their respective number of documents include Li X (90), Wang J (86), Zhang J (84), and Li Y (81). Figure 15 shows the impact ranking of relevant authors according to their H index, a parameter useful in analyzing the performance and impact of an author. The most impactful authors were Zhang J and Liu Y, with H indices of 32 and 31, respectively. Figure 16 showcases the productivity of the top 10 productive authors from 1990 to 2024. Liu Y’s productivity spanned from 1995 to 2024, while Wang J and Liu X started from 2000 and 1999, respectively. The other authors including Wang Y, Zhang Y, Li X, Zhang J, Li Y, Wang X and Li J all had publication start from 2003.

**Figure 14:**
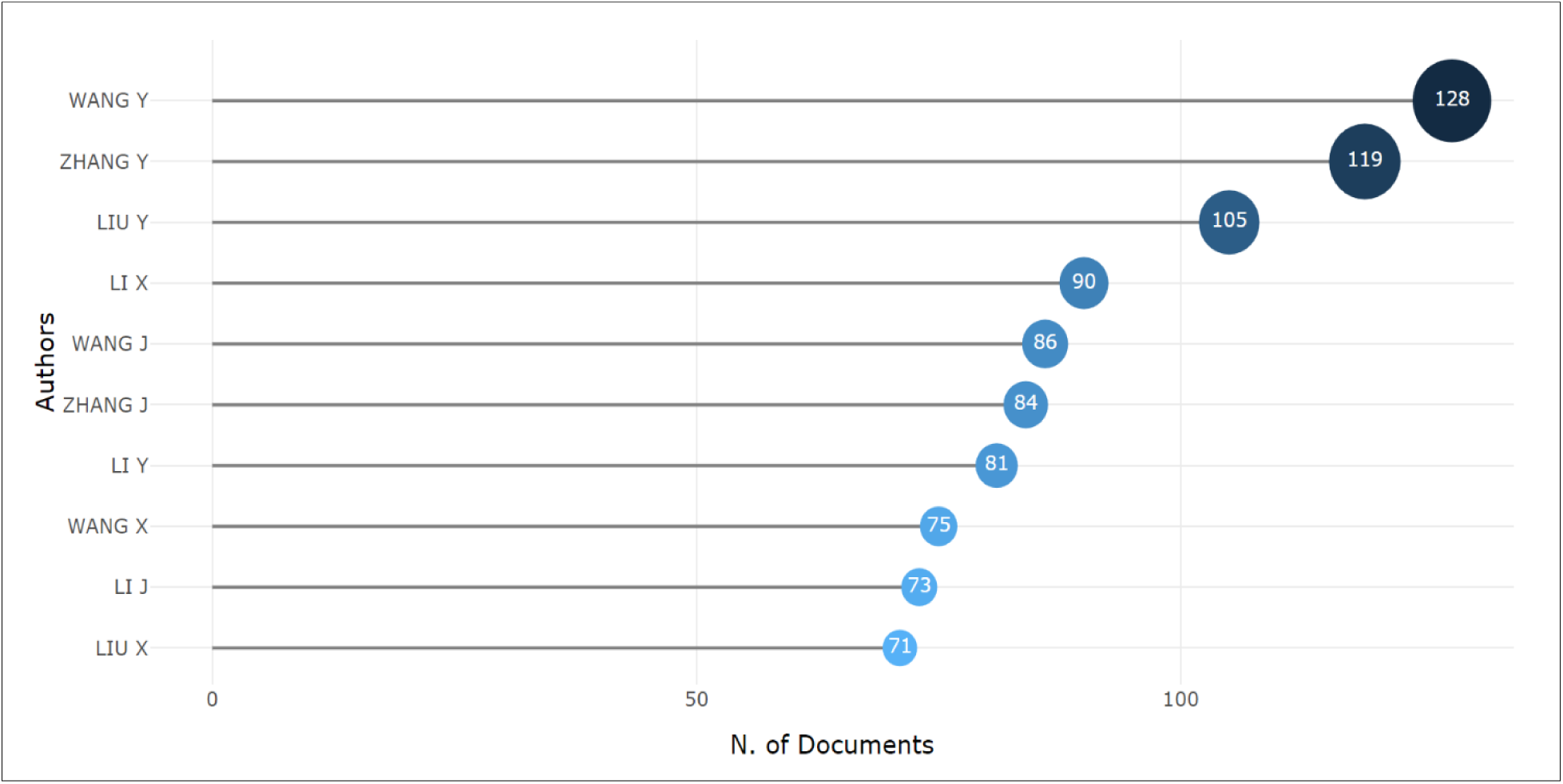
Most relevant authors (based on number of documents) on bioscience research

**Figure 15:**
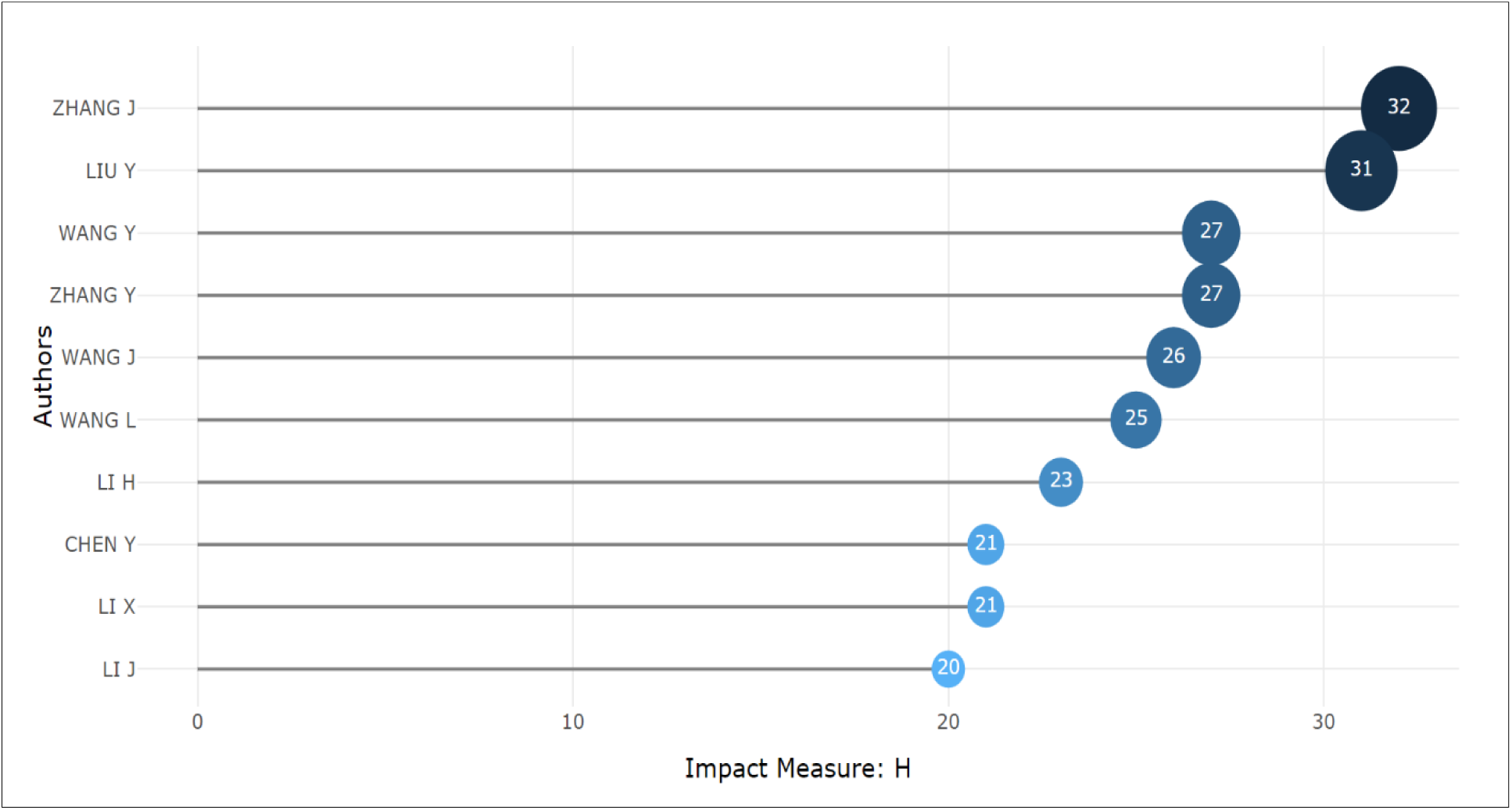
Impact ranking of authors on bioscience research

**Figure 16:**
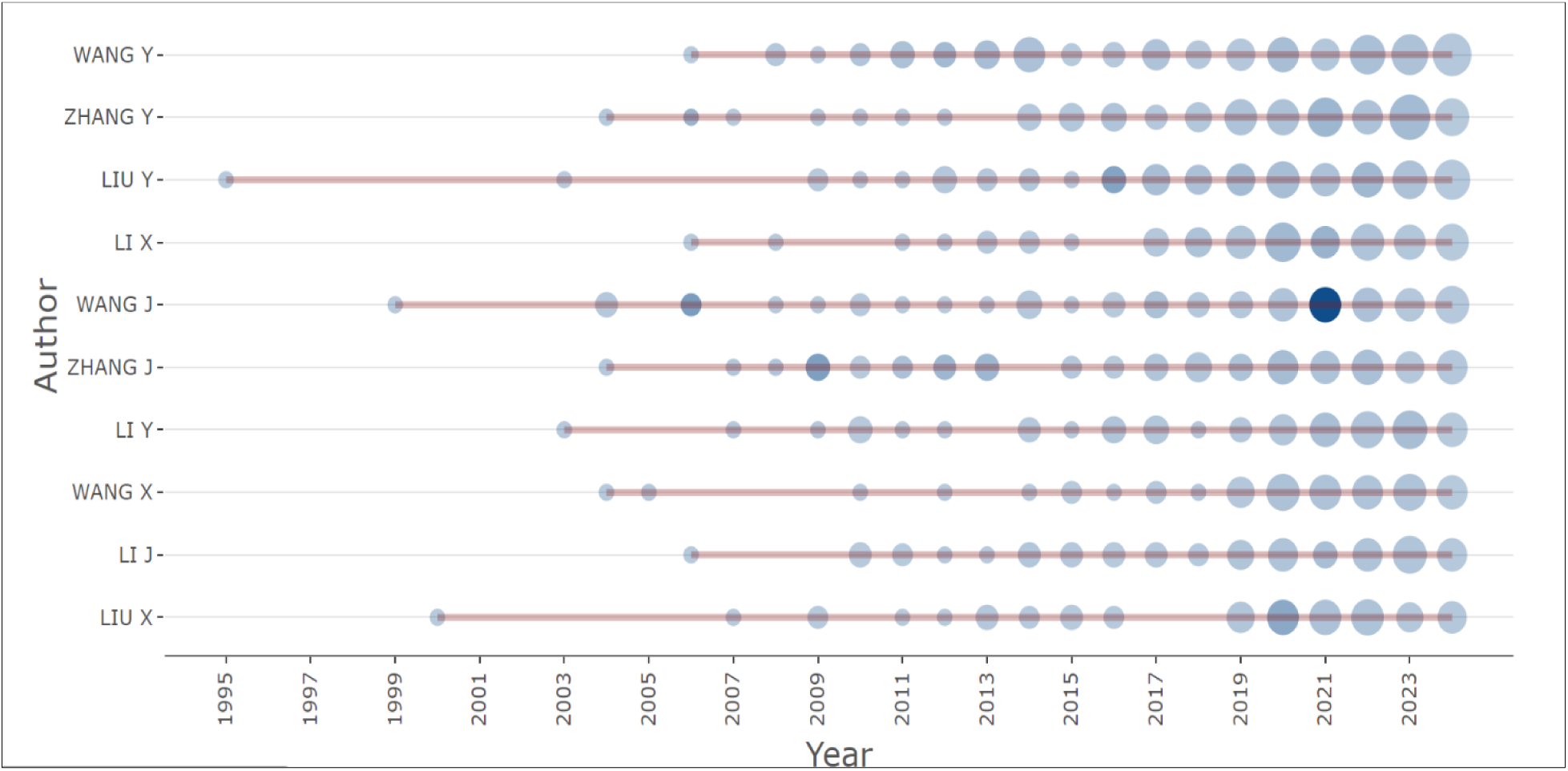
Productivity of Top 10 authors from 1995 to 2024

#### Analysis of keyword evolution

The keywords cluster analysis was done with VOSviewer. The total number of author keywords was 50,012. From these, 5,930 met the minimum threshold of 5 occurrences of a keyword. The overlay visualization of author keyword clusters is shown in Figure 17, while the summary of the most occurring words across the clusters is presented in Figure 18. From Figure 17, concepts such as *methodology*, *rat*, *hybridization*, *histology*, *major clinical study*, and *science* were the dominant themes before 2010. Between 2010 and 2014, the major bioscience research themes were *DNA*, *amino acid sequence*, *animal model*, *cytology*, *education*, and *medical research*. Furthermore, *biological research*, *genetics*, *guide RNA*, *plant genome*, *procedures*, *fluorescence imaging*, *probes*, and *diseases* were reportedly the most occurring keywords from 2015 to 2018. More advanced bioscience themes, such as the *CRISPR/Cas system* and *optical imaging,* showed a high occurrence between 2019 and 2020. As highlighted in Figure 18, the most frequent keywords and trending concepts in bioscience research and education included *biological research*, *genetics*, *procedures*, *metabolism*, *biology*, *DNA*, *gene expression*, *proteins*, *genomics*, and *fluorescence*.

**Figure 17:**
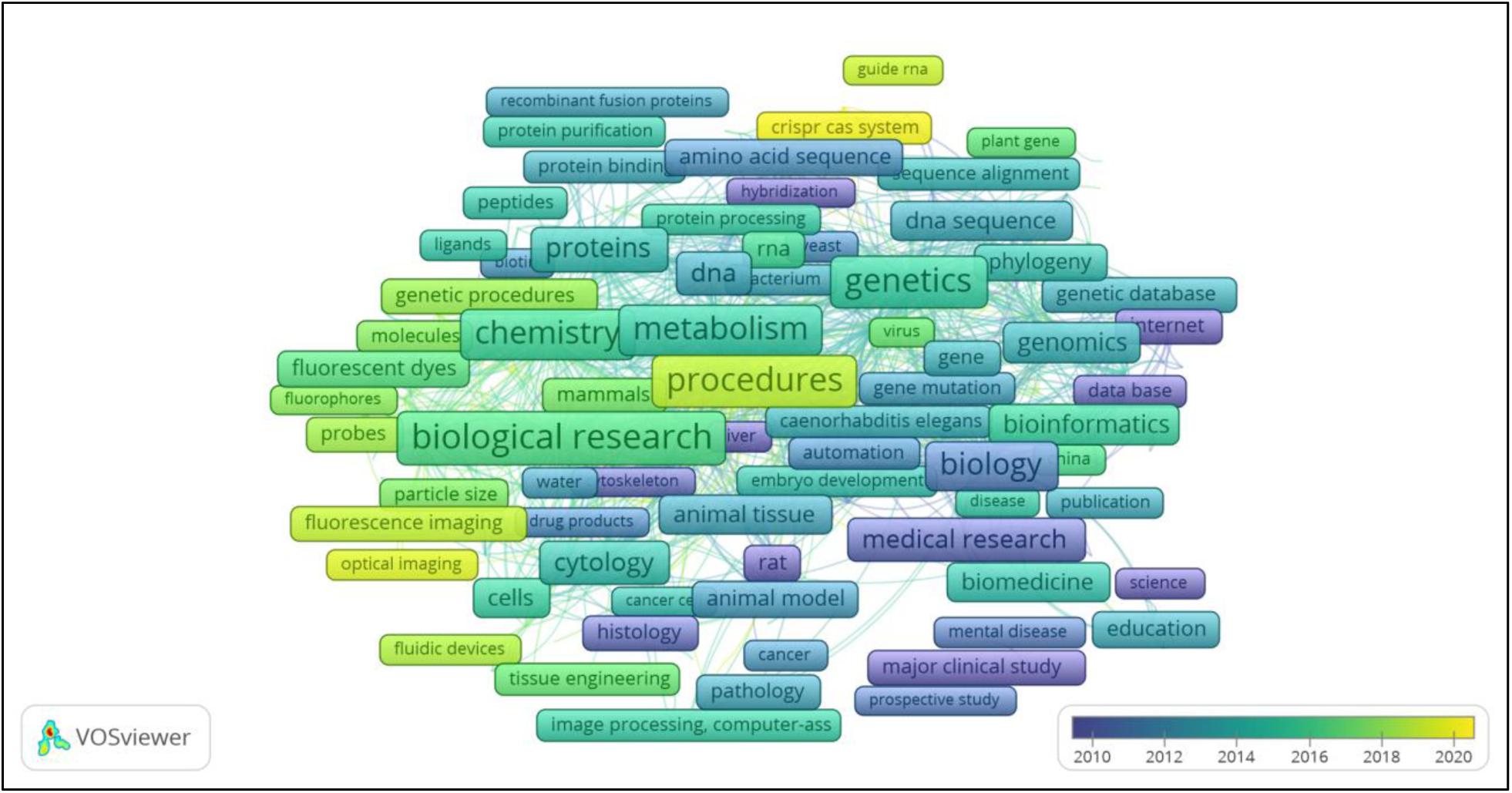
Network of keywords co-occurrence in bioscience research

**Figure 18:**
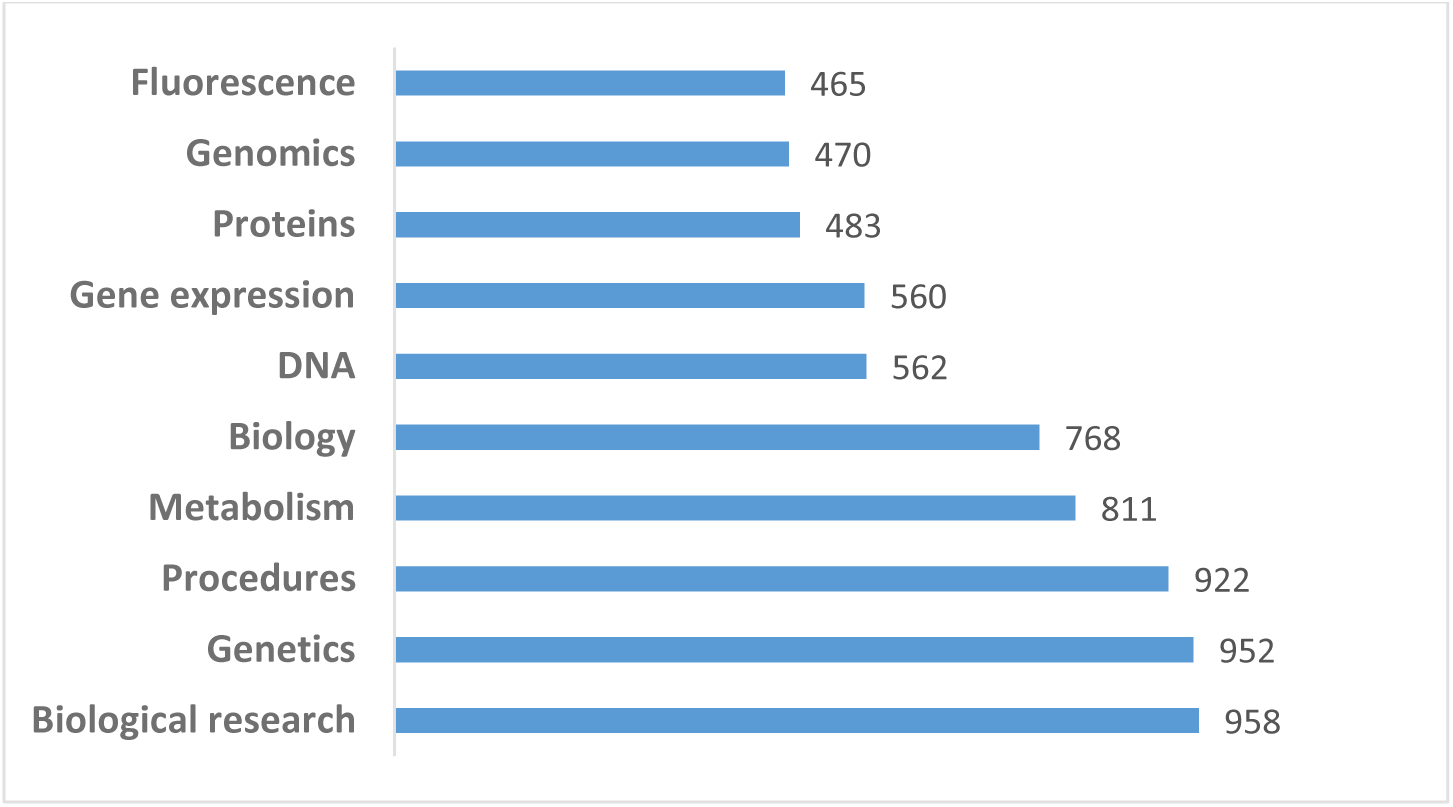
Top 10 most frequent keywords in bioscience research

Out of the 18,589 author-suggested keywords, 704 met the limit of 5 occurrences. From the overlay visualization in Figure 19, *biomass*, *cytology*, *nitric oxide*, *cholesterol*, *insulin*, and *PCR* were depicted in purple and constituted the dominant themes before 2010. Between 2010 and 2018, *bioinformatics*, *fluorescence*, *protein aggregation*, *microfluidics*, *surface plasmon resonance*, *phylogenetics*, *biosecurity*, *microRNA*, *taxonomy*, *epigenetics*, *synthetic biology*, and *sequence analysis* had significant occurrences. Furthermore, the themes shown in yellow, including *chloroplast genome*, *CRISPR/Cas9*, *CRISPR*, *deep learning*, *biological sciences research methodologies*, *biology experiment methods*, *multi-omics*, and *big data,* were the relevant and highly occurring authors’ keywords between 2019 and 2022.

**Figure 19:**
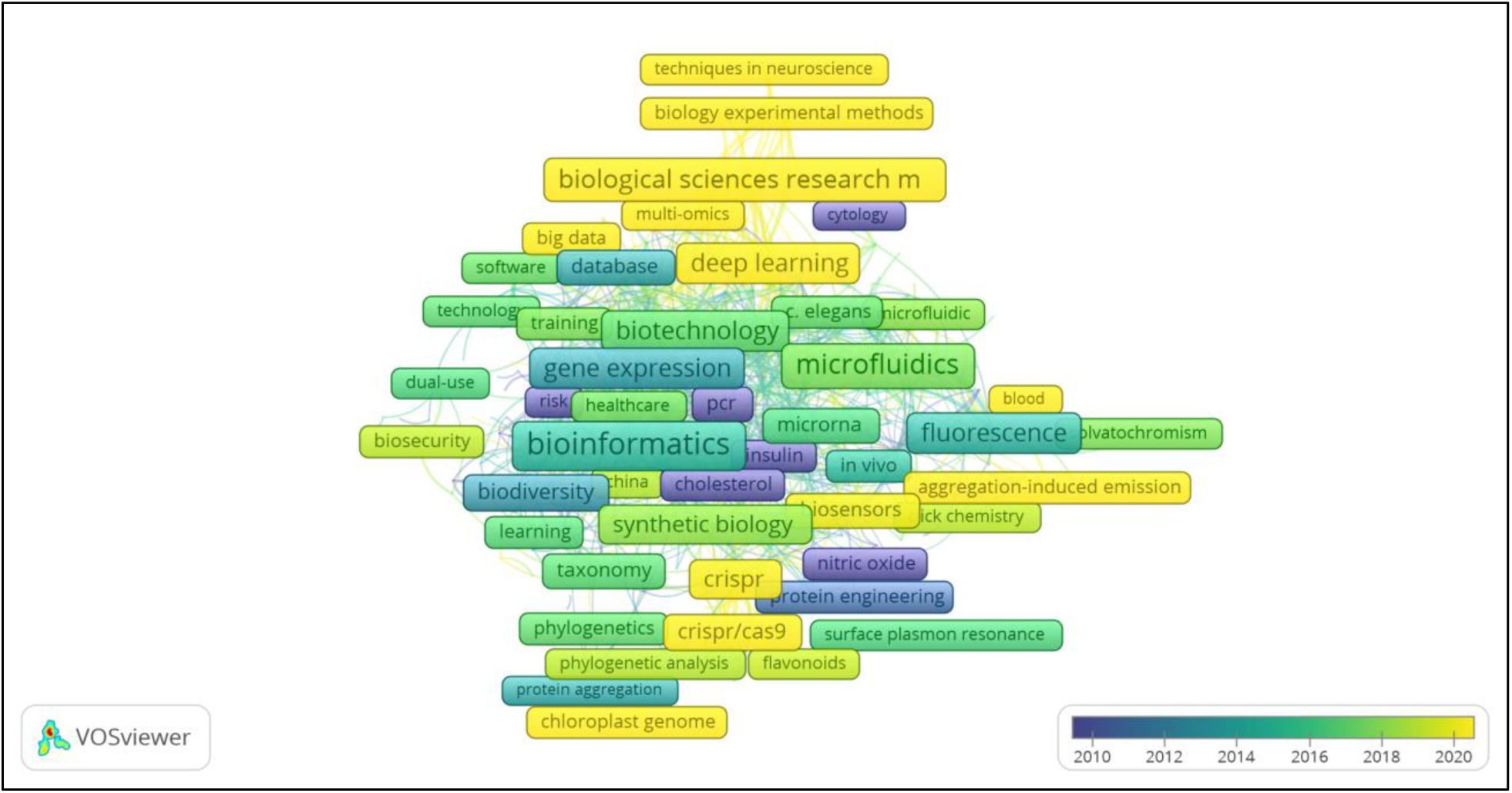
Network of author-suggested keywords in bioscience research

#### Analysis of the most cited documents

The most cited publications on bioscience research are presented in Figure 20 with their details highlighted in Table 2. The majority of the highly cited research papers were review articles. With a total of 23,581 citations, the single-authored research report by Stamatakis (2014) was the most cited. It was followed by Vandesompele et al. (2002) and Hebert et al. (2003), with total citations of 15,901 and 10,446, respectively. The other relevant documents with less than ten thousand citations included Simão et al. (2015), Kilkenny et al. (2010), Steiner (2002), and Beckman and Ames (1998). These documents reflect their importance to the evolving bioscience research landscape.

**Figure 20:**
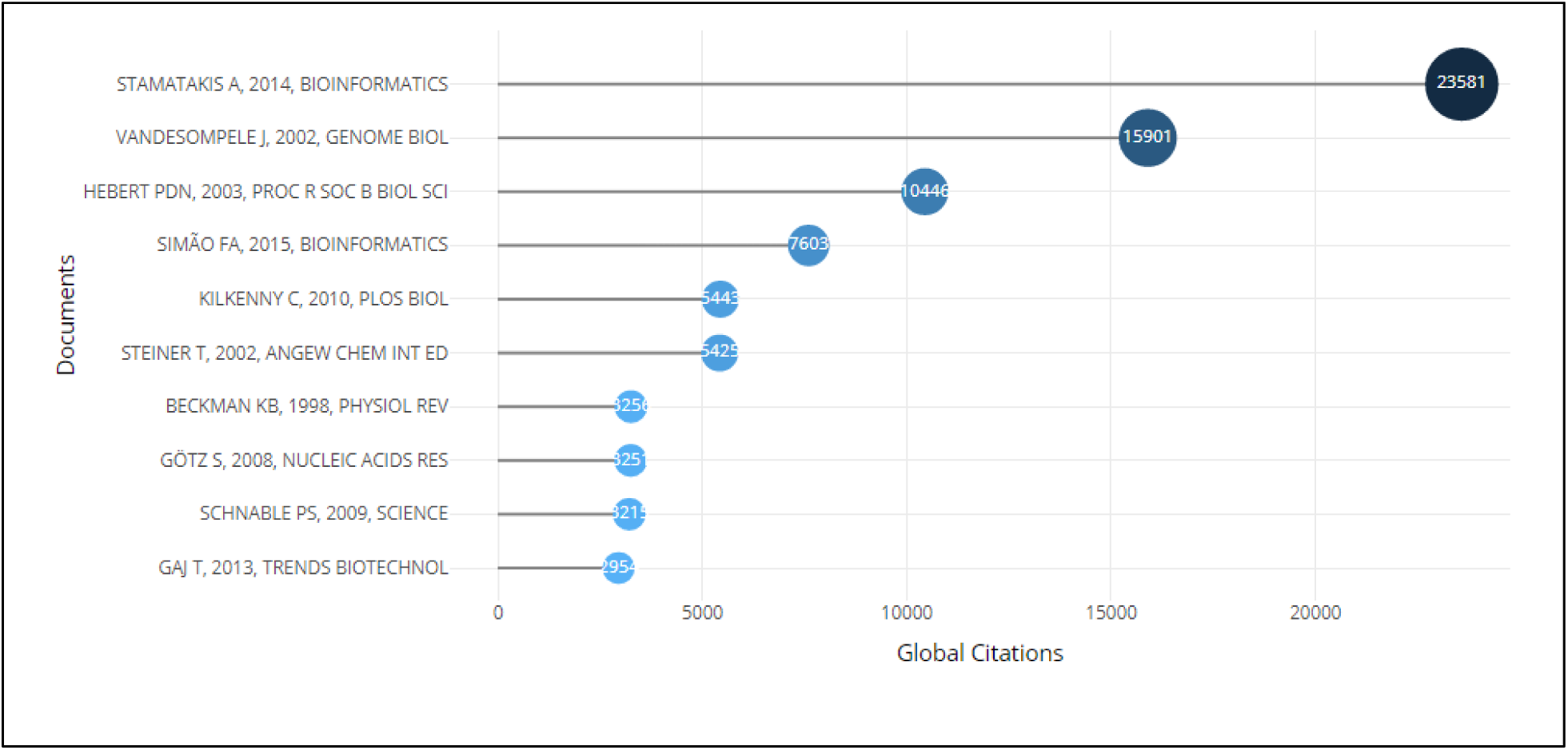
Top 10 most cited documents

## DISCUSSION

### General trends in bioscience research and education

In this study, we used bibliometric techniques to examine the research output and trends in bioscience research and education. There has been a tremendous increase in research output concerning bioscience research and education, indicating that novel concepts, investigations, and deductions are introduced every year, suggesting that the discipline is dynamic. Similarly, the international co-authorships percentage (18.51) as shown in Figure 3 highlights considerable networking in generating scientific output on bioscience. This indicates that international collaboration is responsible for almost one out of every five publications on bioscience research, thus increasing the variety of viewpoints, skills, and funding invested in its research. This draws from the encompassing opportunities of the innovation systems framework and highlights the roles of diverse players in advancing a field.

From the analysis of annual scientific production, it can be deduced that there has been increasing research interest in bioscience, leading to this growth in publications. The highest average total citations of bioscience research and education articles per year were observed in the periods of increasing scientific production. Accordingly, Kelly (2018) highlighted that the bioscience field is publishing more articles in recent years than in the past, which should not be surprising, noting that the volume of publications produced yearly went up by approximately 3.6% between 2000 and 2017, with over 400,000 articles released in 2017.

Due to their wide coverage and impact, research articles and reviews represent the most common means of publishing scientific information. From the present findings, the large number of research articles is a pointer to the nature of scientific publications in biosciences, in which laboratory investigations mostly lead to datasets for original papers. Although it has been acknowledged that research produces a variety of deliverables and implications, Allen et al. (2009) emphasized that the publication of scientific research articles remains a useful indicator of research advancement and knowledge development. Consequently, the large volume of bioscience research output identified in the present analysis suggests that the key actors in the bioscience field are actively involved in the innovations therein. However, the distribution of this advancement across countries presents opportunities for charting a new paradigm for innovations in the subject.

### Country performance and collaborations

A striking finding emerging from the investigation is that the United States and China are leading nations for bioscience research, as reflected in Figures 7A and 7B. However, the presence of the United Kingdom, Japan, Germany, and Brazil as top-performing countries suggests that research on bioscience is localized in the highly industrialized nations. This could be linked to the huge financial and infrastructural investments in research and development in the developed countries. To resolve this imbalance, de Rassenfosse and Selige (2020) suggested that information communication and technology (ICT)–driven research and development knowledge sharing among developed and developing nations must be encouraged, especially in patent transfers, whilst ensuring that developing nations harness their areas of strength for national innovation systems.

Apart from its use in analyzing worldwide networking and knowledge sharing, country co-authorship is also useful in estimating the impact and quality of research ventures. Using the total number of documents computed from VOSviewer, it can be deduced that the United States, China, Germany, Japan, India, and Brazil are key hubs for research linkages due to their strong influence on bioscience research. The numerous links associated with the United States, the United Kingdom, and Germany indicate their high research networking and partnership roles, due to national research policies. The United States has a robust response to bioscience advancement through numerous policies regulating the practice and applications of bioscience-related disciplines, leading to advancements in biotech research protection, and promotion at the state and federal levels (Harris, 2015). The United Kingdom stands as a prominent force in the worldwide race for dominance in the emerging advantages of the biosciences, striving to emerge as an international leader in genomics and bioinformatics (Salter et al., 2016).

The observation that there were more single-country publications than multiple-country publications reflects the need for more inter-country partnerships in bioscience research and education. A country’s achievements in research as well as partnership (local, regional, and worldwide) are related to its economic advancement situation (Payumo and Sutton, 2015). However, the interaction of institutions involved in bioscience, strategic partnerships, and open creativity under a regional and global innovation framework is thought to improve creativity, technology aggregation, and transformative initiatives (Segers, 2016), notwithstanding a country’s development status. This has the potential to strengthen international partnerships.

Furthermore, the results of this study showed that the United States topped the country citations with a total of 96,817 citations. A possible explanation for this might be the relationship between a large volume of publications and higher citations (Larivière and Costas, 2016). Germany, China, and United Kingdom, Belgium, Canada, and Switzerland were part of the most cited countries, conferring on them the leadership status in global bioscience research. The development level of these countries and the huge investments towards research and innovation point to their dominant status in biosciences through increased production of quality, highly cited articles.

### Affiliations, sources, and authors’ contributions

Authors are usually connected to institutions or organizations from which they conduct their research. Thus, affiliations are useful in bibliometric analyses for evaluating research trends and institutional output in a given body of knowledge. Moreover, institutions are an important component of the innovation system. The most relevant affiliations identified from our analysis included the University of California (United States), Universidade de Sao Paulo (Brazil), University of Zurich (Switzerland), Universidade Federal de Sao Paulo (Brazil), Zhejiang University (China), and Tsinghua University (China), and are in the most dominant countries where bioscience research output is significant. The bioscience policies in these nations must have provided an enabling environment for investing into and conducting research in these institutions. This trend shows that these institutions are relevant centers for bioscience research financing, collaboration, and policy making. Opportunities for collaborations, therefore, need to be actively anticipated, especially among researchers from resource-constrained regions (de Rassenfosse and Seliger, 2020).

Scientific information is usually published in sources such as journals and conference proceedings, which are also relevant contributors to the innovative evaluation of research impact. The findings of this study show that the following journals, *Brazilian Journal of Medical and Biological Research*, published by Associação Brasileira de Divulgação Científica (Brazilian Association of Scientific Dissemination), *PLOS ONE*, *Analytical Chemistry*, *Nucleic Acids Research*, *BMC Bioinformatics*, and *Scientific Reports,* published more articles on bioscience. PLOS One, which was established with the intention of quickening the rate of scientific progress and proving its worth, holds that all thorough research should be publicized and should be easily found, broadly circulated, and openly available to everyone (PLOS One, 2025). The findings of cutting-edge studies on the physical, chemical, metabolic, and biological properties of nucleic acids and the associated proteins relating to their metabolic processes, fates as well and reactions are disseminated by *Nucleic Acids Research*, which is published by Oxford University Press (OUP) (OUP, 2024). Furthermore, it is noteworthy that *CBE Life Sciences Education* is one of the influential journals in bioscience, with a particular focus on teaching and learning in the life science field. These scientific journals are devoted to the sharing of knowledge and insights regarding the biosciences.

The present analysis revealed that Scientific Reports, ACS Synthetic Biology, Frontiers in Bioengineering and Biotechnology, Genes, iScience, International Journal of Molecular Sciences, Frontiers in Genetics, and Heliyon showed dominance from 2018 to 2022. This could be linked to the increased interest in omics research, which has great usefulness in health, agriculture, industry, and the environment. For instance, Frontiers in Genetics, which expands knowledge of genomes and gene sequences from humans to plants and other organisms of interest, is regarded as highly cited and draws attention to the latest advancements in genomic techniques, structure, function, and diversity of the genome (Frontiers Media, 2025).

In line with the innovation systems paradigm, authors, by virtue of their expertise, contribute to the innovations in bioscience. A very large number of authors contributing to bioscience research and education (30,380 authors) were reported from the analysis. This shows significant interest and collaboration. However, the majority of the documents were contributions of multiple authors. The reasons for the rise in the number of authors per paper could be that researchers may be discovering new ways to collaborate, combining several areas of specialization to tackle scientific problems, or it could indicate that publishing is becoming more difficult and that joint efforts are needed (Kelly, 2018).

With 128 publications, Wang Y ranked highest according to the number of documents. Also, Zhang Y and Liu Y, Li X, Wang J, Zhang J, and Li Y were key contributors to the share of research documents on bioscience. These authors are linked to China, which is a leading country in bioscience research. Based on the H index, which compares publications and citations, using them to analyze the performance and impact of an author, the most impactful authors were Zhang J and Liu Y with H indices of 32 and 31, respectively. The majority of the authors with a high number of documents were also top-ranking in the H index. Most of the top ranking authors including Wang Y, Zhang Y, Li X, Zhang J, Li Y, Wang X and Li J had publication from 2003, as shown in Figure 16. Furthermore, this analysis highlighted that the publication outputs of the authors peaked from 2020 onwards. This reflects the period of exponential growth in annual scientific production.

### Keywords evolution

Keywords clusters are derived from the keywords identified by the database and the keywords suggested by authors. They contribute to the knowledge base in each field. The findings from the overlay visualization of keyword cluster analysis suggest a transition from traditional biological science research to the recent investigations at the molecular level. Also, the role of precise visualization of biological materials using fluorescent techniques was also evident from the keyword analysis.

From our analysis, the frequently occurring keywords and concepts in bioscience research and education include *biological research*, *genetics*, *procedures*, *metabolism*, *biology*, *DNA*, *gene expression*, *proteins*, *genomics*, and *fluorescence,* as shown in Figure 16. These concepts are the mainstay of research activities and also reflect the current transitions to modern scientific interests in unravelling knowledge at the molecular level. The critical roles of procedures and methodologies in performing bioscience research are also evidenced in this keyword analysis. The capacity to evaluate bioscience investigations or characterize cells and organisms may be limited by the sensitivity of diverse approaches to experimental procedures (Ketchum et al., 2018). The occurrence of *fluorescence* among the most frequent concepts in bioscience highlights its importance as a technique for conducting bioscience research. Originally restricted to cells, fluorescence techniques are now used to analyze real-time protein architecture and behavior in whole animals, providing precise and spatial resolution measurements of biomolecular dynamics (Veerapathiran & Wohland, 2018).

Similarly, the evolution of author-suggested keywords reflects the transition from classical bioscience research themes to recent interests such as *bioinformatics*, *microRNA*, *synthetic biology, CRISPR/Cas9*, *CRISPR*, *deep learning*, *multi-omics*, and *big data*. However, the presence of *biological sciences research methodologies* and *biology experiment methods* indicates the role of experimental procedures in biosciences, as pointed out earlier. The switch in bioscience focus from fundamental studies to the high-impact area of CRISPR/Cas9 reflects landmark moments in technology and financing tactics, particularly in high-income countries. It also shows a move toward practical, translational science with measurable biotechnological, pharmaceutical, and agricultural outputs (Hille et al., 2018). This is now achievable as high-income countries promote technology with economic and health implications, and their quick financial reallocation prioritizes cutting-edge tools (like CRISPR) over basic biology (He and Krainer, 2021).

The key contributions of the most cited documents are summarized as follows. In seeking to develop or combine bio-parts or bio-components for beneficial uses, synthetic biology has recently produced some amazing applications, such as employing light for timed, adjustable, exact cell therapies (optogenetics), bacteria that target tumor cells, cells that rewire human metabolic processes, or engineered live therapeutics which modify the gut-brain-liver axis (Yan et al., 2023). The goal of the established area of bioinformatics is to comprehend biological occurrences by applying statistical techniques and information science, which allows for the collection, examination, and characterization of massive datasets. This aspect of bioscience research is relevant in identifying disease biomarkers and promoting predictive diagnosis. For instance, in order to investigate new biological markers for the prediction of colorectal cancer incidence, Hammad et al. (2021) combined bioinformatics and machine learning techniques. They identified ten (10) genes that were substantially involved in biological events linked to the development of cancer, according to functional enrichment analysis, indicating that these genes with variable expression might be employed as possible biomarkers for the diagnosis of colorectal cancer.

Furthermore, through the use of intricate models made up of numerous layers of nonlinear computing components, deep learning enables the discovery of an arrangement of data with numerous levels of conceptualization (Sapoval et al., 2022). AtomNet, the first structure-based deep convolutional neural network created to forecast small molecule biological function for drug development uses, was presented by Wallach et al. (2022). They gave an example of how to describe biological activity and chemical reactions using the convolutional ideas of feature localization and multilayer structure. The authors claim that AtomNet achieves an area under the curve (AUC) greater than 0.9 on 57.8% of the objects in the DUDE standard, significantly outperforming earlier docking techniques on a wide range of standards.

Using several omics technologies at once presents a previously unknown chance of uncovering molecular characterizations (Zhang et al., 2024). Next-generation sequencing (NGS) advancements have opened the door for new types of omics, including transcriptomics, proteomics, and genomics, metabolomics, with ionomics and phenomics studied among crops (Yang et al., 2021). Moreover, the clustered, regularly interspaced, short palindromic repeats (CRISPR)-associated endonuclease (Cas) system, known as CRISPR/Cas system, has gained enormous attention throughout research sector attributable to its remarkable capacity in turning into diversified treatment interventions in biological and medical domains (Wang et al., 2022). This technique has advanced significantly since its inception and is frequently used in editing genomes to produce point mutations, gene knock-ins, and knock-outs (Tavakoli et al., 2021). As a result, it has been applied in a variety of biological sciences, such as healthcare, plant biology, and livestock breeding.

### Most cited documents

Majority of the highly cited research papers were review articles. In this sub-section, we highlight the key contributions of the top 10 most cited documents. Some of the most noteworthy novel characteristics and upgrades of the Randomized Axelerated Maximum Likelihood (RAxML) were introduced by Stamatakis (2014). These included a significant expansion of alternative models and accepted data types, the addition of SSE3, AVX, and AVX2 vector intrinsics, methods for lowering the code’s memory requirements, and numerous operations for performing post-analyses on arrays of phylogenetic trees. RAxML includes binary, multi-state structural RNA information in addition to proteins and DNA data. For the preparation and analysis of next-generation sequencing data, RAxML provides diverse approaches. To determine whether parts of a gene (such as the 16S) provide a strong and consistent phylogenetic signal that facilitates choices about which parts to multiply, the sliding-window method offered by RAxML holds great promise.

With instantaneous reverse transcription PCR (RT-PCR) emerging as the preferred technique for large-scale and precise expression characterization of specific genes, gene-expression analysis has become more and more significant in biological research. Vandesompele et al. (2002) presented a robust and novel method to estimate the least number of genes needed to compute an objective normalization factor and to select the regulatory genes that are most consistently found in an array of tissues. By analyzing ten housekeeping genes from diverse functional classes and abundances in a range of human tissues, they showed that the traditional approach of using only one gene for standardization results in comparatively substantial mistakes in a sizable percentage of analyzed samples. The authors opined that efficient RT-PCR expression profiling makes it possible to investigate the biological significance of minute expression variations and requires the normalization technique they described.

According to the study of Herbert et al. (2003), one critical way to achieve long-term classification competence is to develop systems that use DNA sequences as taxonomic barcodes. They demonstrated that the cytochrome c oxidase I (COI) gene found in the mitochondria can function as the central component of an international animal bio-identification platform. Following the examination of a single specimen from each of 200 closely related lepidopteran species, a prototype COI characteristic was entirely accurate in recognizing subsequent individuals. The COI identification system offered a dependable, affordable, and easily accessible answer to the existing species identification issue. Significant discoveries about the fundamental principles of evolutionary biology and the diversity of life can be produced by its functionality.

Although genomics has transformed biological research, evaluating the integrity of the entire sequences that are produced is difficult and primarily restricted to technical factors. According to projections for genetic composition shaped by evolution, Simão et al. (2015) suggested a metric for a numerical assessment of genome assembly and identification accuracy. They used sets of Benchmarking Universal Single-Copy Orthologs (BUSCO) to implement the evaluation process in open-source software. For genomes, groups of genes, and transcriptomes, BUSCO quality evaluations offer high-resolution measurements that can be cited in straightforward notation. This makes it easier to measure incremental enhancements to assemblies or to compare newly sequenced genome assemblies with established standards.

Despite the increased the amount of research data that is available in bioscience, a plethora of evidence demonstrates that research reporting practices are frequently insufficient, giving rise to the belief that, even when the science is sound, the articles themselves are frequently miss the intended purposes leading to limited value as tools for shaping scientific policies and practices due to insufficient communication of important details (Kilkenny et al., 2010). Through a team of experts gathered by the authors, the 20 items on the ARRIVE guidelines checklist outline the minimal information that must be included in all scientific reports relating to animal experiments. These include the number and specific features of the animals used (such as species, strain, sex, and genetic background); information about housing and husbandry; and information about the experimental, statistical, and analytical methods (including information about bias-reducing techniques like blinding and randomization). Every item on the checklist has been added to encourage thorough, high-quality reporting so that methods and discoveries may be accurately and critically reviewed.

Steiner (2002)’s review revealed that the most significant directional connection between molecules is the hydrogen bond, and it is used to determine the operations of a wide range of chemical processes, including inorganic and biological, as well as molecular alignment and assembly. Consequently, the study of hydrogen bonding has advanced quickly. The covalent bond, the van der Waals, the ionic, and the cation–π interactions all continuously blend with the hydrogen bond’s wide transition zones. Strong hydrogen bonds can have a very advanced state of proton transfer reactivity, although all hydrogen bonds can be thought of as initiation reactions.

By classifying the literature according to the many kinds of experiments that have been conducted, Beckman and Ames (1998) conducted an in-depth examination of the state of knowledge on the free radical theory. These encompassed epidemiological analyses, human age-related diseases, traditional and population-level molecular genetics, eating habits, metabolic processes, and oxygen concentration, in vitro aging, interspecies analogies, and the continuous clarification of the function of oxygen in biological processes. Although there is increasing agreement that oxidants and oxygen free radicals play a role in progressive aging, scientists have come to the conclusion that there are still many unanswered molecular concerns. This obviously presents research gaps requiring scientific investigations.

In the research of Götz et al. (2008), it was highlighted that the biological study of several species has made extensive use of efficient genomics technology. As details about gene products are frequently essential for interpreting outcomes of experiments, an effective functional annotation of DNA or protein sequences is a crucial prerequisite for the effective utilization of these methodologies. As a result, there is a growing demand for bioinformatics tools that can handle massive volumes of sequence data, generate insightful annotation findings, and are simple for labs working on functional genomics projects to use. Leveraging the Gene Ontology vocabulary, the authors introduced the Blast2GO platform, a complete, biologist-focused solution for the effortless, large-scale functional description of DNA or protein sequences. The combination of several annotation techniques and tools, the many visualization capabilities, the general sequence organizing features, and the high volume capabilities are the most notable aspects of Blast2GO.

The 2.3-gigabase genetic makeup of maize, a significant crop plant and biological research model, has an enhanced draft nucleotide sequence, as described by Schnable et al. (2009) in the genome sequencing of B73 maize. 99.8% of the more than 32,000 estimated genes were positioned on target chromosomes. Numerous categories of transposable elements, distributed unevenly throughout the genome, make up over 85% of the genome. They further described the association of copies of genes with insertions and/or omissions, methylation-poor locations with mutation of the Mu transposon, and the role of unequal gene impairments between redundant regions in restoring an earlier allotetraploid to a genetically diploid condition. Consequently, the cultivation and agricultural advancements of maize are better understood thanks to these evaluations, which also lay the groundwork for future research.

Transcription activator-like effector nucleases (TALENs) and zinc-finger nucleases (ZFNs) are a potent class of instruments that are expanding the frontiers of biological study. These hybrid nucleases consist of a nonspecific DNA-breaking domain connected to adaptable, sequence-dependent DNA-binding subunits. By causing breaks in double-stranded DNA that promote susceptibility to errors, non-homologous end fusion or repair based on homology at particular genomic regions, ZFNs and TALENs allow for a wide variety of genetic changes. Here, Gaj et al. (2013) elucidated the advancements made possible by site-specific nuclease technologies and talked about how these reagents might be used for altering genes and assessment. Furthermore, they illustrated the medical applications of ZFNs and TALENs and suggested the area’s prospective possibilities, including the rise of RNA-guided DNA endonucleases based on the clustered regulatory interspaced short palindromic repeat (CRISPR)/Cas.

From the most cited documents, we deduced that efficient phylogenetics and classification, molecular biology techniques, DNA barcoding and identification, new genomics technologies, and proper research reporting are critical for bioscience research activities. Consequently, the benefits derivable from advancements in bioscience research would largely remain untapped to the extent that these novel techniques are applied across transnational boundaries. As expected, there are ethical issues associated with the practice and techniques of bioscience. Thus, robust considerations for mutually beneficial applications must be encouraged.

### Research and policy issues in bioscience research and education: way forward

This review has identified afresh the inequities in national representation in bioscience advancements. The fundamental roles of new omics techniques and novel concepts such as artificial intelligence, machine learning, and big data in shaping the research landscape on bioscience research and education were part of the findings. Based on the innovation systems framework, a balanced crosstalk among the components is relevant for sustained advancement (Segers et al., 2016). Thus, a global paradigm shift in collaboration and knowledge sharing among developed and developing countries aimed at equity is greatly needed and is hereby advocated for. Key issues for driving this shift are highlighted below:

#### 1. Focus on developing nations

The key actors in bioscience research in developing nations have not featured well in the landscape. To manage societal shifts and support advancements as effectively as possible, localized innovation settings must be fostered and sustained. Such a cluster is made up of local entities and adaptive processes that work together to solve particular challenges. The major components of the ecosystem are top-level universities and research institutes, sufficient investment for new enterprises and research initiatives, a suitable domestic market for novel products, and transnational connections (Oksanen and Hautamäki, 2014).

#### 2. Genomics equity

The poor performance of developing nations in bioscience further buttresses their under-representation in the associated cutting-edge innovations. To promote equity in genomic medicine, steps must be implemented to improve access to genomics research, associated healthcare facilities, and public health initiatives, as well as focus on more research that investigates the therapeutic value of precision therapy among disadvantaged populations (Halbert, 2022). Studies on marginalized populations with a focus on the social and ethical dimensions are needed to address the gaps.

#### 3. Open science initiatives

All the top performing countries in bioscience research had more single-country publications than multiple-country publications. This suggests shortfalls in collaborations. However, the Open Science Framework (OSF) is a project management tool that encourages open, unified operations by allowing the gathering of various components and outputs of research phases, spanning from creating a research hypothesis to conducting a study, preserving and evaluating obtained data, and creating and releasing reports or articles (Foster and Deardorff, 2017). Thus, data access regulations across national borders need to be reviewed to encourage partnerships and ensure open access to repositories.

#### 4. Equitable funds investment

The concentration of bioscience research and innovations in the advanced nations point to prioritization of research funding. High-income countries (HICs) and low-and middle-income countries (LMICs) have influence and resource disparities that prevent many financing strategies from prioritizing the significance of LMIC scientists in influencing global research needs and agendas (Charani et al., 2022). There is also a gap in the money spent by LMIC nations on research concerning their need (Pratt and Hyder, 2018), suggesting a needs assessment. Investors and supporters in developed countries can tackle inequality in their strategies related to research financing through the use of equity funds and discounts for costly consumables.

#### 5. Capacity development

Insights from the present study reveal the low LMIC author share in bioscience research output. This is closely linked with capacity deficits. Large public-academic-private networks can establish novel, fully engaged fellowship initiatives to deal with the substantially small percentage of biomedical professionals in LMICs due to a lack of compatible scientific and professional growth possibilities for early-career researchers (Pillai et al., 2018). Centers of excellence for gene banking and sequencing are practical strategies that can be explored for handling infrastructure deficits.

### Limitations

Despite providing vital information about bioscience research and education studies around the world, this study has some drawbacks. Many other databases, such as Web of Science, Google Scholar, Dimensions, and Lens, were excluded because of the exclusive focus on papers sourced from the Scopus database. This could seem like a disadvantage, but the Scopus database is thought to be the most trustworthy source for scholarly articles. This could, however, result in fewer papers that need to be reviewed. The usage of English-language publications is another restriction. Because of this, the results may be skewed toward authors and sources who speak English and underrepresent other pertinent sources. Leaving out important databases such as Web of Science or PubMed can considerably distort bibliometric reviews, particularly for Latin American and African publications, which are frequently underrepresented in worldwide databases (Asubiaro et al., 2024). About 30-60% of Latin American bioscience papers would be removed from citation analysis due to the under-representation of localized sources, including Scielo and LILACS (Céspedes, 2021). Excluding platforms such as African Journals Online (AJOL) could result in the omission of hundreds of journals, reducing the exposure of regional research (Asubiaro et al., 2024).

Reducing biases and overcoming these constraints should be the goal of future reviews of bioscience research and educational advancements. Despite these limitations, the review’s validity and value remain untouched, offering valuable insights into the current and possible future directions of bioscience research globally.

## Conclusion

The present analysis used bibliometric approaches to examine the general trends in bioscience research and education, country performance and collaborations, affiliations, sources, and author contributions, keywords evolution, and most cited documents, from 1883 to 2024. This paper is the first to evaluate the topic in a long-range perspective using bibliometrics based on the Scopus database, which provides an assessment of research output in the field. The publications analyzed in this study were contributed by 30,380 authors affiliated with diverse institutions across the globe. With an annual rise of 4.41 %, the body of knowledge is growing gradually. Furthermore, the international co-authorships percentage (18.51%) highlights substantial inter-country networking in generating scientific information on bioscience research and education. The United States and China were the most productive and most cited countries, respectively, while the United States had the highest global collaboration. *The Brazilian Journal of Medical and Biological Research* was the most productive source. Wang Y was the most productive author while Zhang J emerged as the most impactful author in the field. *Chloroplast genome, CRISPR/Cas9, CRISPR, deep learning, biological sciences research methodologies, biology experiment methods, multi-omics, and big data* were part of recent concepts that are very relevant in understanding bioscience research and education. Novel concepts such as artificial intelligence, machine learning, and big data have the potential to shape the research landscape on bioscience research and education. This bibliometric study has recognized the current trends that can shape future research on bioscience research and education and has put forward roadmaps for enhancing the proper representation of developing nations in the bioscience landscape.

## Notes

### Competing Interest Statement

The authors have declared no competing interest.

